# Modulation of oscillatory activity in response to very weak intensity transcranial magnetic stimulation in the human primary motor cortex

**DOI:** 10.1101/2025.02.04.636500

**Authors:** Xavier Corominas-Teruel, Martina Bracco, Ann Lohof, Rachel Sherrard, Maria Teresa Colomina, Severine Mahon, Stéphane Charpier, Antoni Valero-Cabré

**Affiliations:** Sorbonne Université, Institut du Cerveau – Paris Brain Institute– ICM, INSERM 1127, CNRS, 7225, APHP-Hôpital de la Pitié Salpêtrière, Causal Dynamics, Plasticity and Rehabilitation Group, FRONTLAB Team, Paris, France; Danish Research Centre for Magnetic Resonance, Centre for Functional and Diagnostic Imaging and Research, Copenhagen University Hospital Hvidovre, Hvidovre, Denmark; Department of Psychology and Research Center for Behaviour Assessment (CRAMC), Universitat Rovira i Virgili, Neurobehavior and Health Research Group, NEUROLAB, Tarragona, Spain; Institut du Cerveau – Paris Brain Institute–ICM, INSERM1127, CNRS7225, APHP-Hôpital de la Pitié Salpêtrière, Sorbonne Université, MOV’IT Team, Paris, France; INSERM1345 & CNRS8263, IBPS-Dev2A, Development, Adaptation and Ageing, Sorbonne Université, Paris, France; Sorbonne Université, Institut du Cerveau – Paris Brain Institute– ICM, INSERM, CNRS, APHP, Hôpital de la Pitié Salpêtrière, Team ‘Network Dynamics and Cellular Excitability’, Paris, France; Cognitive Neuroscience and Information Tech. Research Program, Open University of Catalonia (UOC), Barcelona, Spain; Dept. Anatomy and Neurobiology, Laboratory of Cerebral Dynamics, Boston University School of Medicine, Boston, MA, USA

**Author notes:** Corresponding author: Dr. Antoni Valero-Cabré MD PhD. Cerebral Dynamics, Plasticity and Rehabilitation Group, FRONTLAB team, office 3.028, 48 Boulevard de l’Hôpital, Bâtiment ICM, Paris, France. /.

## Abstract

Transcranial Magnetic Stimulation is widely used to probe and modulate human brain function, yet the neural effects of stimulation delivered at very low intensities remain unclear. Here, we show that very low intensity magnetic pulses can alter ongoing oscillatory activity in the human primary motor cortex. In healthy participants, we combined transcranial magnetic stimulation with electroencephalography to assess neural responses to single pulses and rhythmic stimulation in the motor cortex. Conventional high intensity stimulation produced robust evoked responses and synchronized beta-frequency oscillations. Low-intensity rhythmic stimulation, despite generating much weaker direct responses, modified local oscillatory activity in a manner consistent with phase-dependent enhancement of ongoing rhythms. These findings suggest that cortical oscillations can be influenced by magnetic fields substantially weaker than those typically used in human studies. Low-intensity stimulation may therefore offer a route towards portable, energy-efficient technologies for investigating and modulating brain networks

## INTRODUCTION

Transcranial magnetic stimulation (TMS) at intensities well-below resting Motor Threshold (rMT ^1–5^) has generally been considered insufficient to perturb or modulate neural activity. However, evidence from other neuromodulation approaches strongly suggests that stimulation at weak intensities is able to modulate brain activity. For example, transcranial Electrical Current Stimulation (tCS) employed at intensity levels of ∼0.4 to 0.6V/m cannot directly evoke *per se* measurable physiological effects such as motor evoked potentials or cortically generated phosphene percepts. Yet, it has shown an ability to modulate brain excitability and induce phase entrainment effects in thalamocortical networks by an action on ongoing oscillatory brain intrinsic activity ^6–10^ subtended by ephaptic coupling mechanisms ^11–13^. Likewise*, very low intensity* pulsatile magnetic fields applied to rodent models *in vivo* at peak magnitudes of ∼0.5V/m have recently shown to influence neuronal spiking in locally targeted and interconnected structures ^14,15^. Similarly, recent evidence generated in rats by our own team has shown that a 10-min long 10 Hz repetitive TMS pattern at very low intensity (10mT output) induces a long-lasting hyperpolarization of the resting membrane potential in primary somatosensory (S1) pyramidal neurons, along with sustained reductions in synaptic activity and spontaneous cell firing ^16^. Moreover, rodent studies delivering TMS at intensities well below those conventionally used in humans (0.01-0.07T, ∼0.05-15V/m) have shown a potential to induce cortical plasticity ^17–20^.

Surprisingly, to date, only two studies have systematically assessed the effects of low intensity repetitive TMS in humans. These reports showed respectively that low intensity rhythmic TMS patterns (∼35 or ∼50 V/m) at 10 Hz delivered to the primary visual cortex elicited *online* (i.e., during TMS) signatures of parieto-occipital alpha synchronization ^21^, whereas repetitive rhythmic alpha stimulation (20-pulse bursts repeated 25 times) to the primary motor cortex reduced *offline* (i.e., following rTMS) corticospinal excitability ^22^, hence suggesting that low-intensity magnetic pulsed fields may hold the potential to engage intracortical inhibitory mechanisms in the human motor cortex.

In an attempt to shed further light onto the potential effects of low-intensity TMS, we here aimed to characterize the *online* neural signatures of very weak magnetic pulsed fields in the human brain. To this end, TMS electroencephalography (EEG) ^23^ recordings were employed to assess the effects of ACTIVE single pulse TMS (Sp-TMS) and also rhythmic (high-beta frequency at 20 Hz) TMS bursts (Rhythmic-TMS), delivered to the left primary motor cortex (M1), contrasted to SHAM versions of this same patterns inducing similar scalp/auditory sensations by means of neurally ineffective magnetic pulsed fields. We compared the effects of very weak or low intensity magnetic stimulation (referred in this manuscript as ‘low-intensity’/Li-TMS) delivered at 3% maximal stimulator output, MSO (hence a model predicted E-field strength of ∼6V/m at target) with those achieved with conventional TMS intensity (referred as ‘high-intensity’/Hi-TMS) delivered at 60% MSO (hence an estimated ∼113V/m target E-field strength).

More specifically, aiming to contribute novel and pioneering evidence characterizing their local impact on evoked activity and oscillatory entrainment in the human, we sought to understand if very weak-intensity TMS: (1) perturbed brain oscillatory activity by exploring changes in Event Related Spectral Perturbation (ERSP) and (2) induced local synchronization or entrainment, by a modulation of Inter-Trial Phase Coherence (ITPC); Importantly, we also explored (3)‘state-dependent effects’ exerted by ongoing brain activity at the time of TMS pulse (Fig. 1) onset by characterizing Pre-TMS oscillatory phase-dependent effects on TMS elicited ERSPs. We observed that both single TMS pulses and 4-pulse rhythmic TMS bursts delivered at a high-beta (20 Hz) frequency at high intensity (Hi-TMS) entrain beta-band oscillations in the left primary motor cortex. Most importantly, low-intensity (Li-TMS) rhythmic TMS also modulates oscillatory activity in this same region, with effects suggesting a phase-dependent modulation of ongoing local rhythms. Our findings support the ability of very weak intensity TMS to modulate cortical oscillatory activity in M1 in a brain state-dependent manner, while also highlighting the need for further mechanistic research and for online and offline characterization of the dose response relationship of TMS delivered at weak intensities.

**Figure 1.**
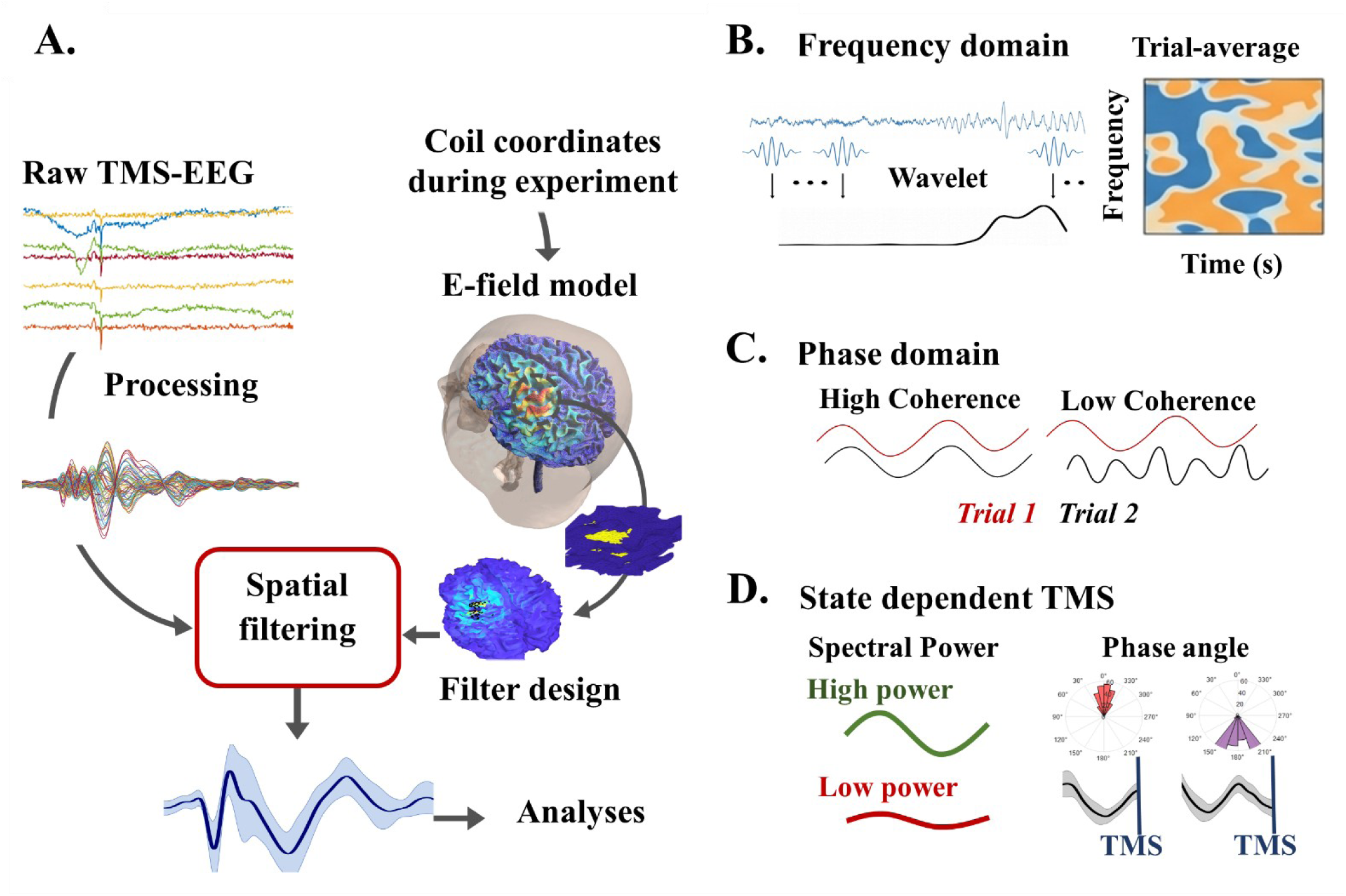
TMS-EEG data analysis pipeline employed in our study. **(A)** MRI-Neuronavigated TMS-EEG was used to characterize the *online* impact (i.e., during stimulation periods) of *Single Pulse* and *Rhythmic* (20Hz frequency) TMS patterns on the left M1 cortex of our cohort of healthy participants. To this end, EEG data was pre-processed using a pipeline able to identify and subtract artifactual signals. In parallel, individualized tetrahedral finite element head models (FEM) reconstructed from T1w MRI volumes for each participant were used to estimate the E-field distribution and magnitude delivered during experimental sessions to the left M1 cortex hotspot for the right First Dorsal Interosseous (FDI) hand muscle. This head model served to estimate the magnetic field strength and to delineate for each participant a specific region of interest (ROI) defined as the 80^th^ percentile of the predicted E-field strength. Minimizing mass conduction and multisensory afferent signal components, a spatial filtering pipeline encompassing the targeted regions of each specific participant was applied to reconstruct TMS modulated EEG sources and extract EEG activity signatures elicited by stimulation from the primary targeted region (EPICURUS signal processing pipeline). Finally, analyses in the **(B)** time-frequency and **(C)** phase-domains were conducted to explore TMS spectral perturbations (ERSP), TMS inter-trial phase synchronization (ITPC) and oscillatory **(D)** phase-dependent (bound to brain state) modulatory effects on ERSP.

## RESULTS

### Analysis of Event-related spectral perturbations (ERSP)

We first tested whether TMS at high and very weak intensities was able to perturb ongoing brain oscillations by means of a cluster-based analysis in the time and the frequency domains comparing the time-frequency power spectra of local M1 cortical activity. As expected, Event-related Spectral Perturbations (ERSP) cluster-based statistics performed on ACTIVE *vs*. SHAM single-pulse TMS at conventional high-intensity (Hi-TMS, 60%MSO) yielded a cluster of early elicited spectral power increase (∼0-100 ms, p<0.0004, *d*=0.7) encompassing the beta band ([10-35 Hz]), while a comparison of ACTIVE vs. SHAM high-intensity rhythmic TMS bursts revealed a longer event of beta oscillatory activity (∼150-300 ms, p<0.0004, *d*=0.8) outlasting the TMS burst itself (from 0 to 150ms; Fig. 2). On the other hand, cluster-based permutation analysis failed to reveal significant differences in time-frequency power for the ACTIVE *vs*. SHAM single pulse condition or for the low-intensity rhythmic TMS condition (Li-TMS, 3% MSO), as suggested by the lack of statistically significant clusters (p<0.4) across the examined frequency bands ([8-35 Hz]) and time windows ([0-600 ms]) (see Fig. 2, bottom row).

**Figure 2.**
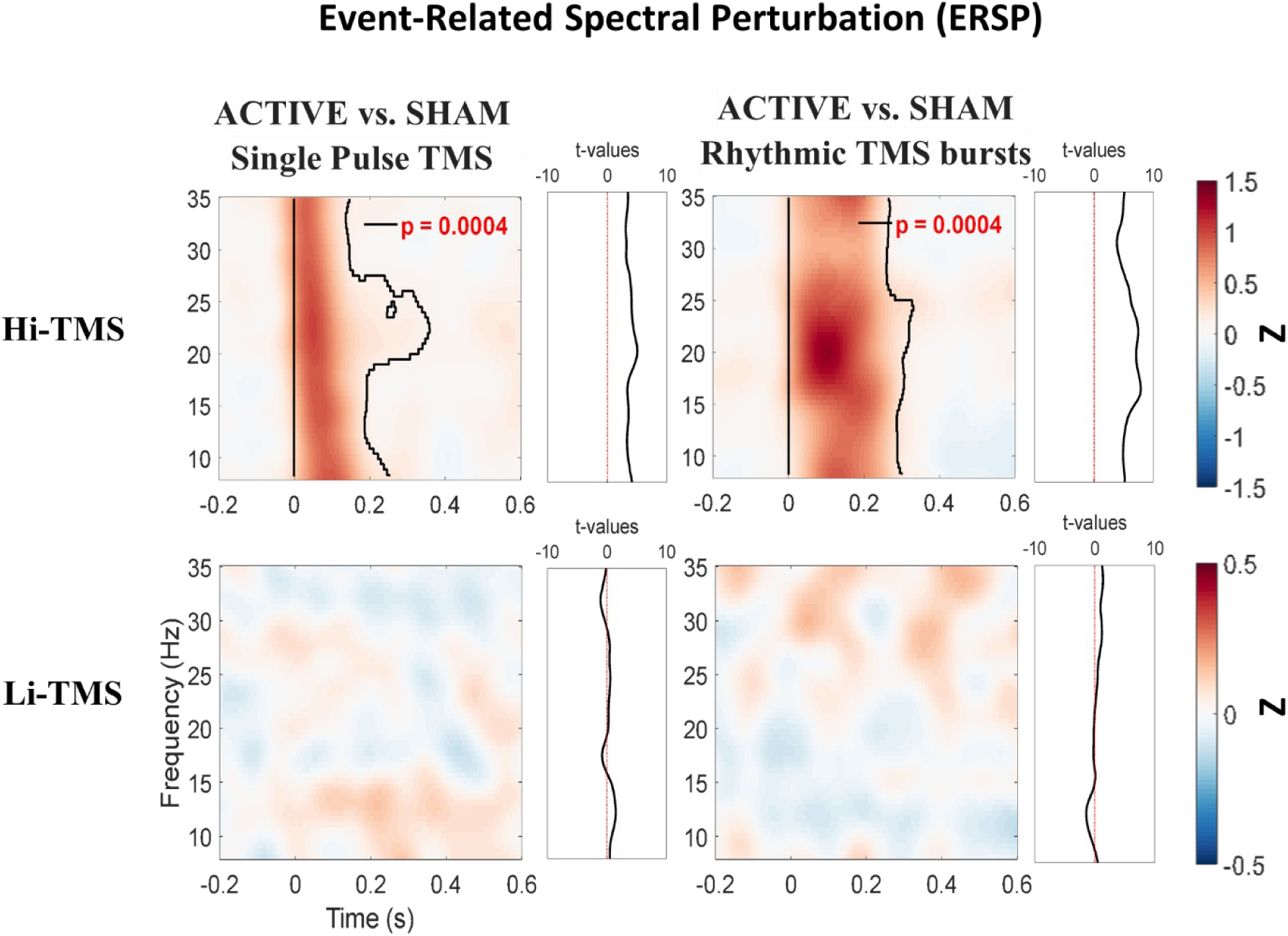
Effects of high and very weak-intensity TMS on Event-Related Spectral Perturbation (ERSP) measures. Group-level (n=18) Time-frequency maps displaying ERSP across experimental conditions. All trials have been averaged across participants and grouped by specific TMS conditions (*Single Pulse TMS* and *Rhythmic TMS* at high and very weak-intensity stimulation, referred as Hi-TMS and Li-TMS respectively). ERSP data displayed in the figure correspond to Z-transformed group grand averages organized for each TMS condition. Comparisons between ACTIVE vs. SHAM independently for *Single Pulse* and *Rhythmic* TMS and for high-intensity (Hi-TMS) and very weak-intensity (Li-TMS) TMS pulses or bursts are presented in the figure. Black outlines indicate statistically significant EEG clusters in the time and frequency domains (p<0.05, two-sided). Hi-TMS: High-Intensity TMS (60%MSO, ∼113V/m, cohort mean relative ∼78.9%rMT), Li-TMS: very weak-intensity (3%MSO, ∼6V/m, cohort mean relative ∼3.8%rMT). rMT: resting Motor Threshold; MSO: Maximal Stimulator output.

### Analysis of inter-trial phase coherence (ITPC) and frequency-specific entrainment

We then investigated whether TMS at high and very weak intensities induced phase resetting and entrainment of brain oscillations through a cluster-based analysis in the time-frequency domain comparing local M1 inter-trial phase coherence (ITPC). ITPC cluster-based statistics performed on ACTIVE vs. SHAM single-pulse TMS at high-intensity (Hi-TMS, 60%MSO) revealed a significant cluster of early (∼0-300 ms, p<0.0001, *d*=0.6) alpha to high-beta (8-35 Hz) phase synchronization across trials, with a pronounced enhancement near 30 Hz. A contrast between ACTIVE vs. SHAM high-intensity rhythmic TMS bursts yielded a significant cluster of early (∼0-300 ms, p<0.0001, *d*=0.7) alpha-to-high-beta (8-35 Hz) phase-synchronization across trials, this time with a pronounced high-beta enhancement around 20 Hz resulting in an episode of beta synchronization outlasting the duration TMS bursts (from 0 to 150 ms). In contrast, no significant ITPC clusters (p<0.3) were observed for any of the low-intensity TMS conditions (Li-TMS, 3% MSO) across the time and frequencies tested, neither for ACTIVE vs. SHAM single-pulse TMS nor for ACTIVE vs. SHAM rhythmic TMS comparisons (Fig. 3).

**Figure 3.**
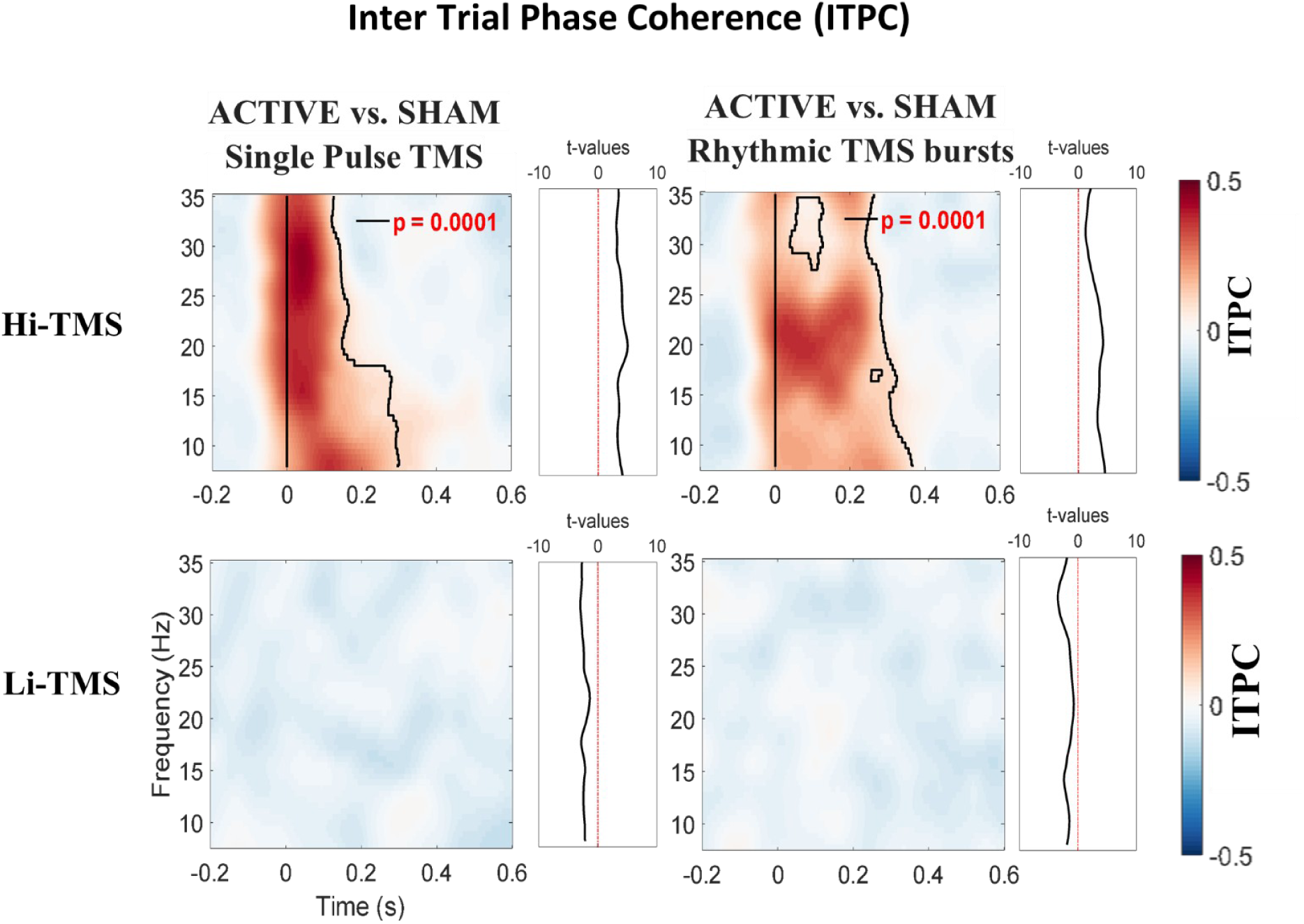
Effects of high and very weak-intensity TMS on Intertrial Phase Coherence (ITPC) measures. Group-level (n=18) Time-frequency maps displaying ITPC within the motor cortex region across experimental conditions. Displayed ITPC data correspond to grand averages across participants grouped by each TMS conditions tested in our study (*Single Pulse TMS* and *Rhythmic TMS* at high and very weak-intensity stimulation, Hi-TMS and Li-TMS respectively). Comparisons are shown for ACTIVE vs SHAM for *Single Pulse TMS* and *Rhythmic TMS* (after subtracting their respective SHAM conditions) for high-intensity (Hi-TMS) and very weak-intensity (Li-TMS) stimulation TMS pulses or bursts. Black outlines indicate statistically significant EEG clusters in the time and frequency domains (p<0.05, two-sided).

### Analysis of Alpha and Beta pre-TMS phase-dependent effects

To evaluate phase-dependent ERSP and ITPC, individual EEG trials were categorized according to their pre-TMS alpha phase ([8-13 Hz]) and pre-TMS beta phase ([14-30 Hz]). These two frequency bands were specifically chosen based on prior empirical evidence indicating that alpha activity in pre-central regions, largely driven by sensory areas, reflects strong feedforward inhibition between S1 and M1, whereas beta activity is the dominant oscillatory pattern associated with motor cortex activation ^24–26^.

### Alpha pre-TMS phase-dependent effects

To specifically assess whether spectral perturbations were modulated by the oscillatory state of ongoing brain activity at the time of stimulation, hence show ‘state-dependency’ effects, defined by pre-TMS mu-alpha oscillatory phases (considered a proxy of intrinsic fluctuations of excitability in M1), we statistically compared measures of ERSP power for mu-alpha phase-sorted trials, contrasting values at the *peak vs*. the *trough* of the ongoing oscillation prior to ACTIVE TMS (after subtracting their respective SHAM TMS condition).

ERSP cluster-based analysis did not reveal significant time-frequency clusters (p<0.2) for single-pulse TMS delivered at either high intensity (Hi-TMS, 60% MSO) or low intensity TMS (Li-TMS, 3% MSO). Statistics performed on rhythmic TMS at high-intensity yielded early (∼15 Hz, 100-350 ms, p<0.03, *d*=0.5) and late (25 Hz, 200-400 ms, p<0.03, *d*=0.3) clusters in the tested time-frequency domain with increased spectral power when the first pulse of the TMS burst was aligned with the *peak* of the ongoing mu-alpha oscillations. Additionally, a third early cluster emerged at higher frequencies (∼35 Hz, 0-100 ms, p<0.03, *d*=-0.4) suggesting increased spectral power when the onset of the first pulse of the TMS burst was aligned with the *trough* of the ongoing mu-alpha oscillations. Finally, statistics performed for low-intensity rhythmic TMS conditions yielded an early cluster (∼20-25 Hz, 0-100 ms, p<0.04, *d*=-0.2) suggesting increased spectral power when the onset of the first pulse of the TMS burst was aligned with the *trough* of the ongoing mu-alpha oscillations (Fig. 4).

**Figure 4.**
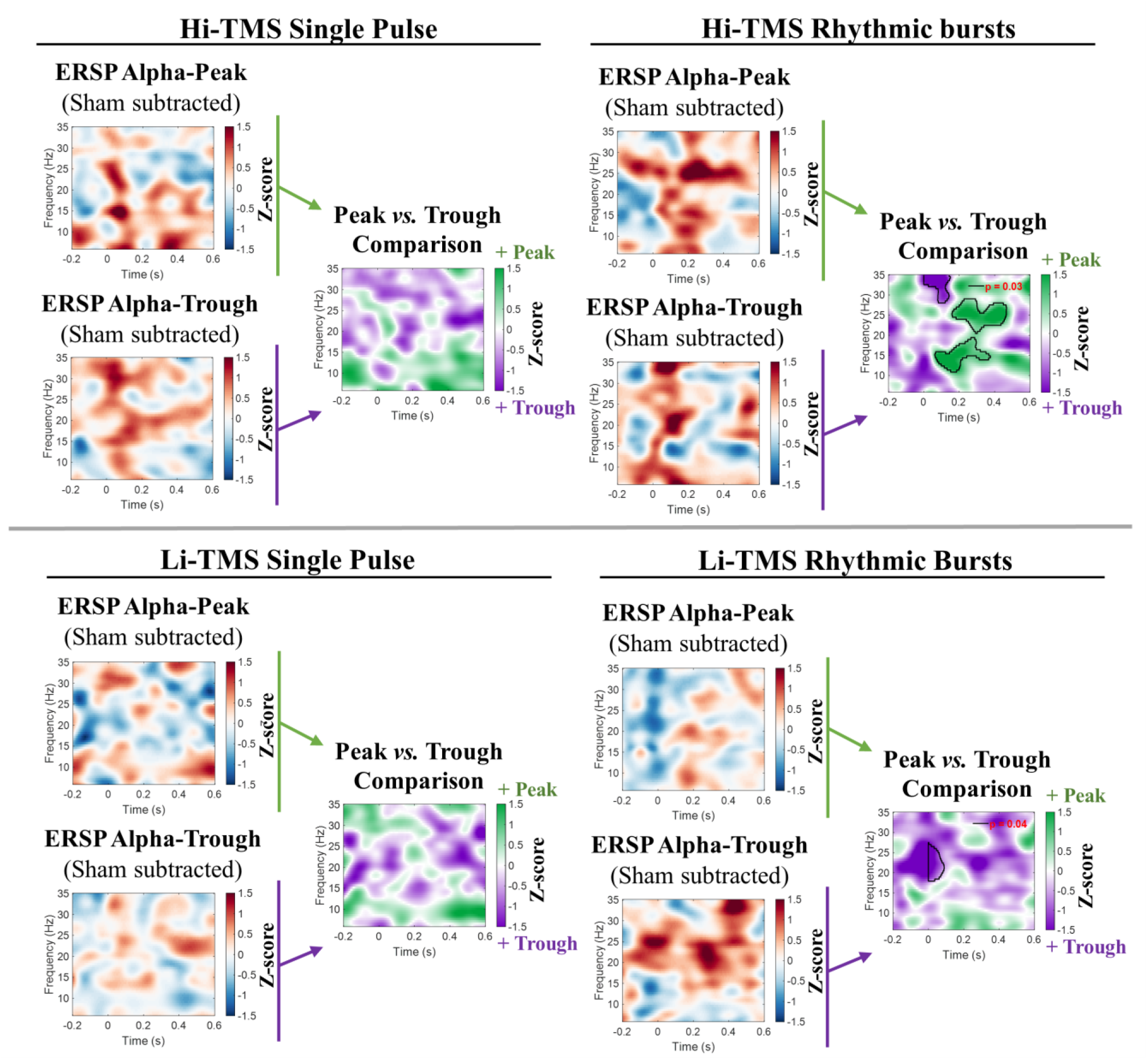
Alpha phase-dependent TMS effects on Event-Related Spectral Perturbation (ERSP) estimates. Group-level (n=18) Event-Related Spectral Perturbation measurements sorted according to the instantaneous pre-TMS onset phase (-5ms to -2ms) of the mu-alpha rhythm (8-13Hz). ACTIVE TMS trials (exceeding +50% of total oscillatory power (-400ms to -3ms) across all ACTIVE trials, SHAM subtracted) at the *peak* or the *trough* of the alpha rhythm at the time of 1st TMS pulse onset were identified and selected for final analysis. Per every tested TMS condition, the phase-dependent time-frequency Z-score (relative to their respective minimum and maximum spectral power) for ACTIVE (SHAM subtracted) is independently displayed, where ***Red areas*** indicate oscillatory power increase, and ***Blue areas*** indicate oscillatory power decrease. Moreover, the comparisons between TMS conditions contrasting *peak* vs *trough* sorted trials are displayed in the figure. ***Green clusters*** of EEG plots indicate oscillatory power increase for the ERSP when TMS pulses were delivered around the *peak* of the ongoing oscillations. ***Purple clusters*** of EEG plots indicate oscillatory power increase from the ERSP when TMS pulses were delivered around the *trough* of the ongoing oscillations. Black outlines indicate statistically significant clusters between conditions in the time and frequency domains (p<0.05, two-sided). Hi-TMS: High-Intensity TMS (60%MSO, ∼113V/m, cohort mean relative ∼78.9%rMT), Li-TMS: very weak-intensity (3%MSO, ∼6V/m, cohort mean relative ∼3.8%rMT). rMT: resting Motor Threshold; MSO: Maximal Stimulator output.

In an attempt to further explore the nature of the above-reported alpha phase-dependent influences, ACTIVE and SHAM ERSPs were statistically compared *post-hoc* only for ACTIVE (SHAM-subtracted) conditions in which differences between the ERSP *peak* and *trough* had previously reached statistical significance. Statistical analyses of rhythmic TMS for conventional high intensity TMS (Hi-TMS), comparing ACTVE and SHAM pulses delivered at the alpha-*peak* phase revealed an early significant cluster (∼25 Hz, 150-450 ms, p=0.01, *d*=0.6), of increased spectral power during ACTIVE stimulation. Similarly, analyses for low intensity TMS (Li-TMS), comparing ACTIVE and SHAM pulses delivered at the alpha-trough phase revealed both, early (∼23 Hz, 0-400 ms, p<0.02, *d*=0.3) and late (∼33 Hz, 400-450 ms, p<0.02, *d*=0.3) significant clusters, also indicating increased spectral power during ACTIVE stimulation (see Supplementary Figs. 1 & 2).

Finally, to assess whether inter-trial phase coherence was influenced by brain state as defined by the phase of pre-TMS mu-alpha ongoing oscillations, we also performed statistical comparisons on ITPC for mu-alpha phase-sorted trials with the same approach previously followed for ERSPs. Specifically, we contrasted values at the *peak* vs. the *trough* of the ongoing oscillation for ACTIVE TMS conditions (after subtracting their respective SHAM patterns, see Supplementary Fig. 3). Statistical comparisons of single-pulse TMS at conventional high intensity (Hi-TMS) revealed a late significant cluster (∼10 Hz, 400-500 ms, p=0.003, *d*=0.5) suggesting higher levels of late-phase synchronization across trials when TMS was delivered at the peak of the ongoing alpha oscillation. To further characterize alpha phase-dependent ITPC interactions, ACTIVE and SHAM high intensity single-pulse ITPC conditions were post-hoc compared (see Supplementary Fig. 4) revealing an early significant cluster (∼30 Hz, 150-450 ms, p=0.01, *d*=0.6) for pulses delivered at the alpha-*peak* phase, hence indicating increased ITPC during ACTIVE TMS stimulation. Note that no additional significant clusters or trends were observed for any of the other tested conditions.

### Beta pre-TMS phase-dependent effects

Paralleling analysis performed for alpha pre-TMS phase-dependent effects, we also tested whether spectral perturbations showed state-dependent behavior, according this time to pre-TMS beta oscillatory phase. We hence computed parallel statistical comparisons for ERSP spectral power based on beta phase-sorted trials (*peak* vs. the *trough*) for the ACTIVE TMS condition (after subtraction of their respective SHAM conditions). ERSP cluster-based analysis performed on single-pulse TMS at conventional high-intensity revealed an early (∼15 Hz, 0-100 ms, p<0.03, *d*=-0.5) cluster in the time and frequency domain with increased spectral power when the onset of stimulation was aligned with the *trough* of the ongoing beta oscillations. Additional significant high-frequency clusters (∼33 Hz, 200-400 ms, p<0.03, *d*=-0.4) suggested increased spectral power when the TMS pulse onset was aligned with the *peak* of the ongoing beta oscillations. ERSP cluster-based analysis performed on single-pulse TMS this time at low intensity revealed a non-significant trend (∼10 Hz, 0-200 ms, p<0.07) of increased spectral power when the TMS pulse onset was aligned with the *trough* of the ongoing beta oscillations (Fig. 5).

**Figure 5.**
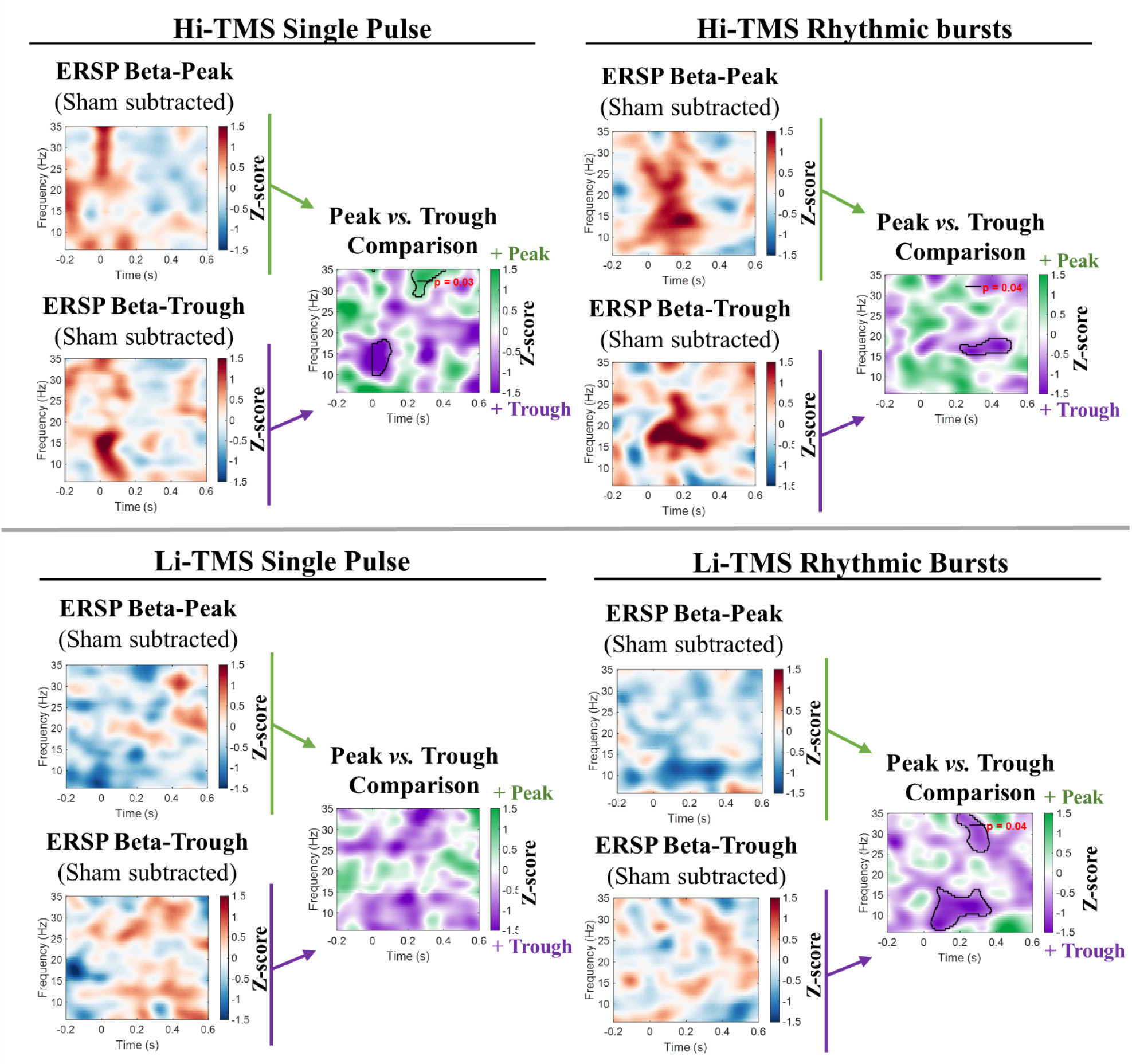
Beta phase-dependent TMS effects on Event-Related Spectral Perturbation (ERSP) estimates. Group-level (n=18) ERSP measurements sorted according to the instantaneous pre-TMS onset phase (-5ms to -2ms) of the beta rhythm (14-30Hz). ACTIVE TMS trials (exceeding +50% of total oscillatory power (-400ms to -3ms) across all ACTIVE trials, SHAM subtracted) at the *peak* or the *trough* of beta rhythm at the time of the 1st TMS pulse onset were identified and selected for final analysis. For each tested TMS condition, the phase-dependent time-frequency Z-score (relative to their respective minimum and maximum spectral power) for the ACTIVE (after SHAM subtraction) condition is independently displayed. ***Red-colored areas*** indicate oscillatory power increases, whereas ***Blue-colored areas*** indicate oscillatory power decreases. Moreover, the comparisons between TMS conditions contrasting *peak* vs *trough* sorted trials are displayed in the figure. ***Green-colored clusters*** of EEG plots indicate oscillatory power increases for the ERSP when TMS pulses were delivered around the *peak* of the ongoing oscillation. ***Purple-colored clusters*** of EEG plots indicate oscillatory power increase from the ERSP when TMS pulses were delivered around the *trough* of the ongoing oscillations. Black outlines indicate statistically significant clusters between the mentioned phase conditions in the time and the frequency domains (p<0.05, two-sided). Hi-TMS: High-Intensity TMS (60%MSO, ∼113V/m, cohort mean relative ∼78.9%rMT), Li-TMS: very low intensity (3%MSO, ∼6V/m, cohort mean relative ∼3.8%rMT). rMT: resting Motor Threshold; MSO: Maximum Stimulation Output.

Statistical comparisons performed on rhythmic TMS at high intensity yielded late clusters (∼15-20 Hz, 200-500 ms, p<0.04, *d*=-0.5) with increased spectral power when the onset of first pulse of the TMS burst was aligned with the *trough* of the ongoing beta oscillations. In contrast, only non-significant trends were observed for early clusters (∼10 Hz, 0 ms-200 ms, p<0.06/∼25 Hz, 0-100 ms, p<0.06) when the first pulse of the TMS burst was aligned with the *peak* of the ongoing beta oscillations.

Finally, statistical comparisons performed for low-intensity rhythmic TMS patterns yielded an early significant cluster (10-15 Hz, 0-400 ms, p<0.04, *d*=-0.1) outlasting the TMS burst (from 0 to 150 ms), and a late significant cluster (30-35 Hz, 300-400 ms, p<0.04) both showing increased spectral power when the onset of the first pulse of the TMS burst was aligned with the *trough* of the ongoing beta oscillations (Fig. 5).

To further explore the nature of beta phase-dependent interactions, ACTIVE vs. SHAM TMS-modulated ERSPs were statistically compared *post hoc* only for conditions in which ERSP *peak* vs. the *trough* differences (SHAM-subtracted) had reached statistical significance. Statistical analysis of single-pulse TMS at conventional high intensity (Hi-TMS), comparing ACTIVE vs. SHAM conditions for pulses delivered at the beta-*trough* phase, revealed both, early (∼15 Hz, 0-150 ms, p<0.05, *d*=0.6) and late (∼35 Hz, 250-450 ms, p=0.006, *d*=0.4) significant clusters, indicating respectively, early increases and late decreases in spectral power during ACTIVE stimulation (see Supplementary Figs. 5, 6 & 7). Similar analyses for conventional high-intensity rhythmic TMS showed that pulses delivered at the beta-*peak* phase yielded an early significant cluster (∼10-20 Hz, 0-250 ms, p<0.05, *d*=0.4), suggesting increased spectral power during ACTIVE stimulation. Similarly, pulses delivered at the beta-*trough* phase revealed an early significant cluster (∼15-20 Hz, 0-400 ms, p<0.001, *d*=0.4), also indicating increased spectral power during ACTIVE stimulation. In contrast, low intensity rhythmic TMS bursts (Li-TMS) with pulses delivered at the beta-*peak* phase, yielded an early significant cluster (∼10 Hz, 0-400 ms, p=0.01, *d*=-0.2), suggesting this time decreased spectral power during ACTIVE TMS.

Finally, to assess whether inter-trial phase coherence effects were also influenced by brain state defined by pre-TMS beta oscillatory phase, statistical comparisons on ITPC were performed for beta-phase-sorted trials. Specifically, ITPC time-frequency values at the *peak vs*. the *trough* of the ongoing oscillation were contrasted for the ACTIVE TMS condition (after subtracting their respective SHAM patterns). No significant clusters or trend-level effects were observed in any of the tested conditions (see Supplementary Fig. 3).

## DISCUSSION

In the present study, we employed concurrent TMS-EEG to assess the potential impact of very weak pulsatile TMS fields (referred in the manuscript as Low Intensity, Li-TMS, 3% MSO, ∼6V/m) on the human brain. Equivalent conventional Intensity patterns (referred in the manuscript as High intensity, Hi-TMS ∼60% MSO, ∼70-140V/m) served as a physiological benchmark for interpreting the effects of the former.

Replicating outcomes from prior studies, and as expected, single pulse M1 high intensity TMS yielded transcranial evoked potentials (TEPs; data unreported in the main text, see Supplementary Fig. 8 for full TEPs time-series) and prototypical ERSP and ITPC ^26–33^, while rhythmic high intensity TMS bursts at 20 Hz elicited *online* time- and frequency-dependent spectral perturbation and entrainment phenomena, as shown by changes in ERSP and ITPC (see Figs 2 & 3, respectively). Prior research has shown that spontaneous fluctuations in neocortical rhythms regulate in a phase-dependent manner, the timing of information flow ^34–36^. In the present study, high-beta (20 Hz) rhythmic TMS administered at conventional high intensity (Hi-TMS), although below the individual resting motor threshold, elicited stronger beta oscillatory activity when the onset of the rhythmic TMS bursts was aligned with the peak of ongoing alpha oscillations. This suggests that rhythmic TMS may be more effective at inducing entrainment when cortical activity is inhibited, as lower endogenous fluctuations enhance sensitivity to externally imposed rhythms. In contrast, brain regions exhibiting strong oscillatory reverberation appear to demonstrate greater resistance to externally imposed rhythmic stimulation and reduced susceptibility to phase-resetting effects ^37,38^. Accordingly, sorting TMS trials by beta rhythm phase revealed that high-intensity TMS delivered at the *trough* of the ongoing beta oscillation resulted in increased low-beta-imposed oscillatory activity, suggesting that this phase corresponds to a cortical state with lower endogenous resistance, thereby facilitating phase-resetting. However, since rhythmic TMS bursts and single pulse TMS at high intensity are likely subtended by different physiological mechanisms, further research, outside the scope of the present work, should explicitly confirm and investigate the basis of such an unexpected observation.

Prior work in rodents using electrical stimulation has shown that weak currents may selectively modulate alpha rhythms, originating prominently from the spiking of inhibitory interneurons ^10^ or superficial agranular cortical layers. Likewise, human neuroimaging and pharmacological studies have reported a causal role for alpha oscillations increasing local inhibitory tone ^25,39–44^. Prior studies have also unearthed that weak (30V/m E-field strength) and mild intensity (50V/m E-field strength) TMS might be sufficient to entrain cortical alpha oscillatory activity and to modulate corticospinal excitability ^21,22^. Although our study employed lower intensities (Li-TMS; 5-12V/m induced current) and a distinct stimulation frequency (20 Hz), comparable outcomes were observed. Notably, beta-band oscillatory activity was increased when low-intensity rhythmic TMS pulses were delivered during locally disinhibited oscillatory states, represented by the trough of mu-alpha oscillations. This facilitatory effect is consistent with prior findings suggesting that the trough of alpha oscillations enhances neural input recruitment ^26,37^, thereby promoting, when paired with low intensity TMS, oscillatory activity at the primary motor cortex natural frequency. Moreover, beta oscillations in the motor cortex are thought to be primarily generated by recurrent excitatory-inhibitory interactions within local cortical circuits, involving particularly pyramidal neurons and fast-spiking GABAergic interneurons (PV+) in superficial layers ^25,42–44^. In that context, the observed beta enhancement may reflect recruitment of these local circuits, whereby disinhibited states (e.g., alpha *trough*-phase) increase input gains and allow rhythmic low-intensity TMS to entrain activity at the system’s natural resonance frequency. Our results also revealed a decrease in alpha activity when ACTIVE rhythmic low-intensity TMS was delivered at the peak of ongoing beta oscillations. In line with our own findings, specific phases of the beta cycle might reflect fluctuations in cortical gain shaped by underlying excitatory-inhibitory dynamics. Hence the delivery of stimulation during such low-gain states may bias local circuitry toward an inhibitory state, limiting effective neuronal recruitment and disrupting pulse-facilitation-inhibition processes subtended by alpha rhythms across sensorimotor networks.

Several limitations of the present study should be acknowledged. First, despite randomized experimental conditions and measures to mitigate auditory/somatosensory confounds, residual differences between Hi-TMS and Li-TMS (via somatosensory input and bone-conducted effects) could have persisted. Even if comparable across conditions, the effect of Hi-TMS inherently elicited larger-amplitude responses than Li-TMS, limiting the validity of direct comparisons. Accordingly, qualitative comparisons regarding the nature of the effects allow more appropriate and meaningful interpretations and were privileged. Second, although intensity for Hi-TMS (60% MSO) remained below motor threshold levels for all participants (group mean: 78.6% of rMT), indirect influences via somatosensory feedback or subtle movement-related artifacts on EEG signal cannot be fully excluded. Third, even if randomly interleaved to maximize signal-to-noise ratio for phase-dependent effects, the number of ACTIVE TMS trials was set to be twice that of the SHAM TMS ones, to allow a robust estimation of state-dependent modulations of oscillatory activity, while maintaining a sufficient number of SHAM trials to control for non-specific effects. However, to mitigate statistical biases, post hoc ACTIVE *vs*. SHAM analyses were performed using equalized trial counts, with ACTIVE trials randomly subsampled to match the number of SHAM trials. Fourth and last, our study employed fixed TMS intensities (Hi-TMS at 60% MSO and LI-TMS 3% MSO) across participants, even if MRI based modeling (SimNIBS toolbox) individually predicted the magnitude of the induced electric field (E-Field, in V/m) on each participant, as an approximation of field strength at target. However, E-field predictions strongly depend on biophysical assumptions limiting the interpretation of the MSO-based intensity levels delivered on M1, an important consideration for future low-intensity TMS research.

Overall, extending prior evidence, the current study presents to the best of our knowledge pioneering evidence suggesting that the modulatory effects of very weak TMS intensity here described are not fully frequency dependent. This outcome is likely explained by the fact that stimulation at such a low intensity might not be strong enough to impose an external rhythm. Instead, very weak intensity TMS at levels here tested, may facilitate the spontaneous increase or entrainment of endogenous activity, mostly affecting neural populations on the superior cortical layers. Importantly, within sensorimotor systems, inhibitory interneurons known for possessing low firing resistance ^10,42,45^ could be predominantly perturbed, which may explain and generate the phase-dependent rhythm effects observed in our study. Moreover, the fact that rhythmic TMS bursts at very weak-intensity synchronized activity in the beta range and desynchronized activity in the alpha range, whereas the same patterns delivered at high-intensity facilitated activity in the beta band, indicates that divergent mechanisms are engaged by the two stimulation protocols, solely differing (same stimulated region, with identical procedures and control conditions) in their intensity levels. On such a basis, while high-intensity TMS might entrain frequency-specific oscillations, we here suggest very weak-intensity TMS may act by amplifying in a brain-state dependent manner spontaneous activity from highly sensitive intraneuronal populations of the sensorimotor cortex. Further studies employing rodent single unit electrophysiology and computational E-field predictions coupled to realistic cortical neuronal models will be required to identify the predominant cell-type(s) and mechanisms involved in these phenomena. In parallel, further human trials assessing the *online* and *offline* influences of TMS bursts, repetitive TMS patterns, and dose-response relationships between elicited oscillatory activity will help pinpoint the future potential of very weak-intensity TMS, while paving the way for the development of portable, battery-operated multichannel low-intensity TMS devices able to operate on local and extended brain networks in future exploratory and clinical avenues.

## METHODS

### Participants

We recruited eighteen healthy volunteers (n=18, mean age ±SD = 26.1 ± 5.8 years; 10 females / 8 males), all right-handed according to a standard handedness inventory test ^46^, free of neurological, or psychiatric symptoms and with normal or corrected-to-normal vision. All participants satisfied international TMS and magnetic resonance imaging (MRI) safety criteria ^47,48^ and provided informed consent prior to the study. All ethical regulations relevant to human research participants were followed. The research protocol respected the Declaration of Helsinki, French legislation regulating good clinical practices (ICH-E6, R2) and received the approval of a local ethical committee (*Comité de Protection des Personnes, Ile de France I*). No participant reported serious discomfort or adverse events to TMS ^49^.

### Experimental design

Participants took part in two different sessions. In the first one, a 3D-T1-weighted MRI structural scan (3T Siemens MP2RAGE, flip angle = 9, TR = 2300 ms, TE = 4.18 ms, slice thickness = 1 mm, isovoxel) was acquired. In the second, TMS was applied to the participants’ left M1, while 64-electrode scalp EEG was concurrently recorded. The primary motor cortex was selected as a target region due to its well-characterized anatomy and the extensive body of TMS-EEG literature, providing a solid framework for interpreting our findings ^50–54^. In addition, the M1 region allows stimulation effects to be referenced to relative individual excitability measures, namely the resting motor threshold, rather than relying solely on absolute TMS Machine Stimulation Output (%MSO) values. Participants were exposed to four stimulation conditions: *Single-pulse* TMS (Sp-TMS), and short *rhythmic* 20Hz TMS (Rhythmic TMS), each delivered at low intensity (*Li-TMS*) and at conventional High-Intensity (*Hi-TMS*; see TMS section for further details). As the effects of each TMS condition were evaluated in its ACTIVE and SHAM modalities, each participant was assessed on a total of 4 ACTIVE TMS conditions (and their four associated SHAM equivalents). TMS conditions were randomly counterbalanced across participants to avoid order bias effects ^55^.

### Transcranial Magnetic Stimulation (TMS)

Stimulation was applied over the left M1 with a 70 mm figure-of-eight coil (D70 alpha (a) coil) connected to a biphasic rTMS stimulator (Magstim Rapid2, UK). The left M1 was identified as the optimal cortical hotspot on which TMS pulses elicited motor evoked potentials (MEPs) over the right first dorsal interosseous (FDI) muscle. Prior to the start of the experimental session, we measured for each participant the rMT, by delivering single pulses of TMS over the M1 while recording MEPs amplitudes (Signal 5.0, Cambridge Electronic Design Limited, Cambridge, UK). Starting at 65%MSO, TMS was delivered at varying intervals between ∼6-8 secs and adjusted with intensity increases/decreases at 1-5% MSO steps, according to the amplitude and consistency of the evoked MEPs responses. The individual rMT was defined as the lowest TMS intensity able to induce at least 5 out of 10 MEP (50% responses) with a minimal peak-to-peak amplitude of 50μV (group rMT average: 77.2 ± 7.1 %MSO; range: 68-95 %MSO; Note that the rMT was individually estimated with the EEG grid in place, hence it resulted in higher values than those usually measured directly by posing the TMS coil on the scalp). During the whole session, TMS coil position was tracked using an MRI-based frameless stereotactic neuronavigation system (Brainsight, Rogue Research) and kept within a 2 mm radius from the center of the established targeted M1 hotspot site.

For conventional high-intensity blocks (*Hi-TMS*), TMS intensity was set at a fixed level of 60% of the MSO (68.8 dI/dtmax [A/μs]) ^56^; corresponding to a relative 78.6 ± 6.5% of rMT; range: 63.2-88.2% rMT; Supplementary Table 1, for individual rMT details). For low (very weak) intensity TMS blocks (*Li-TMS*), we established an intensity of 3% of the MSO (3.44 dI/dtmax [A/μs]; corresponding to: 3.86 ± 0.34% of rMT (range: 3.1-4.4% rMT), which was the lowest TMS intensity level eliciting consistent pulse output in our TMS equipment (see Supplementary Note 1 for details). This intensity was selected as the closest approximation of low-intensity TMS stimulation levels previously employed with a miniaturized custom-made device in a rodent study ^16^ according to the constraints of a commercially available human TMS equipment (Magstim Rapid2 and a D-70 mm figure-of-eight coil; see Supplementary Note 1). Each of the 4 experimental conditions (Hi-Sp TMS, Li-Sp TMS, Hi-Rhythmic TMS, Li-Rhythmic TMS), included the administration of two blocks of 60 trials. Rhythmic TMS consisted of trains of 4 pulses delivered at 20 Hz (inter-stimulus interval (ISI) of 50ms with a total burst duration of 150 ms). Each block encompassed 40 ACTIVE single TMS pulses and 40 ACTIVE TMS bursts and 20 randomly-embedded SHAM pulses and bursts, resulting in a total of 120 trials per participant and condition, with an inter-trial interval (ITI) of 6 secs (hence 80 ACTIVE and 40 SHAM pulses/bursts per participant) for each of the conditions. SHAM pulses/bursts were delivered by a second rTMS device (Magstim Super Rapid^2^) using an identical figure-of-eight TMS coil (D70 mm double coil) oriented perpendicularly and placed right above the ACTIVE coil, so that the magnetic field was projected away from the scalp.

### TMS-EEG data acquisition

EEG was continuously recorded with a TMS-compatible EEG amplifier (actiChamp, Brain Products) attached to a 64 active electrode grid (ActiCap snap), mounted on an elastic cap following International 10-10 EEG system positions. FPz and Fz served as *ground* and *reference*, respectively. Signals were sampled at 25 kHz and impedances were periodically monitored and kept below 5kΩ. To minimize TMS-related auditory evoked EEG responses, participants wore foam earplugs through which a white masking noise was continuously played (ER-3C insert earphones, ETYMOTIC Research) at a volume tailored individually to ensure participants were unable to notice the TMS “clicking” sound when pulses were delivered. To minimize muscle and eye movement artifacts, participants were seated in a comfortable armchair with their heads on a chinrest and required to remain relaxed with their gaze fixed on a cross (1 cm x 1 cm) displayed in the center of a computer screen placed at 57 cm from the participant.

### TMS-EEG pre-processing

Signal preprocessing was performed with MATLAB (The MathWorks Inc., Natick, MA, USA) using EEGLAB software (UC San Diego, La Jolla California, USA; ^57^) and its TMS-EEG signal analyzer plugin (TESA; ^58^). Every patient’s dataset was preprocessed individually according to the following steps: First, EEG data was epoched around the TMS pulse onset (or the 1^st^ pulse of each burst, [-1.200 ms,1.200 ms]). TMS-pulse electromagnetic artifacts were removed and zero-padded ([-2 ms,10 ms]) and the recharging artifact removed and interpolated using a spline cubic function ^59^. Following, datasets were downsampled to 5 KHz, and the most artifacted channels and trials were identified and temporarily excluded. Eye movements and continuous muscle artifacts were removed through Independent Component Analysis (ICA; ^60^), while the SOUND algorithm was used to suppress recording noise, and the SSP-SIR filter to remove time-locked TMS-evoked muscle artifacts ^61,62^ if present. Datasets were bandpass filtered with a fourth-order Butterworth filter ([2-85 Hz]), downsampled to 1KHz, baseline corrected and re-referenced to an average of all channels. Finally, to improve the signal-to-noise ratio, we implemented an E-field based spatial filter employing the EPICURUS method (E-field based sPatial fIltering for Concurrent EEG-TMS data Used to taRget soUrce Signal) developed in our laboratory (see ^63^ and Supplementary Note 2 for details). Briefly, a source-based spatial filter based on the individual distribution of the TMS coil magnetic field was developed for each participant (see Supplementary Fig. 9 for participant-specific E-field models and spatial filter designs). The spatial filter was employed to highlight the signal from the primary stimulated location (M1) while minimizing distant signal crosstalk leakage from non-directly stimulated brain regions increasing signal to noise ratio without manipulating unfiltered sensor-level ERSP or ITPC dynamics (see Supplementary Figs. 10 & 11) ^64^.

### Statistics and Reproducibility

Postprocessed signals were analyzed using MATLAB, and the MATLAB-based Fieldtrip toolbox ^65^. Our analyses sought to understand if very weak-intensity stimulation was able to (1) perturb brain oscillations, (2) induce local synchronization or entrainment, and (3) understand the brain state-dependent effects (based on EEG phase dependency). Accordingly, the following metrics were respectively analyzed each of these three objectives: (1) Modulation of Event-Related Spectral Perturbation (ERSP); (2) Modulation of Inter-Trial Phase Coherence (ITPC); and (3) Pre-TMS oscillatory phase-dependent effects on elicited ERSP.

For spectral perturbation analyses (ERSP), signals were time-frequency transformed ([8-35 Hz]) in steps of 0.01Hz using complex Morlet wavelets ^65–67^ and Z-transformed concerning the mean and standard deviation of the entire epoch length ([-0.5 s to 0.6 s]). For local entrainment analyses (i.e., ITPC), phase synchronization across trials was estimated immediately after the time-frequency transformation. Finally, to evaluate phase-dependent ERSP and ITPC, individual EEG trials were categorized according to their pre-TMS alpha phase ([8-13 Hz]) and pre-TMS beta phase ([14-30 Hz]). As indicated previously, our analytical focus on these two frequencies is based on prior empirical evidence supporting a role for alpha pre-central activity in feedforward S1 to M1 inhibition, and beta activity as dominant oscillatory pattern associated with motor cortex activation. To this end, a first window of interest preceding the TMS pulse (1^st^ TMS pulse for rhythmic burst conditions) was isolated ([-5 ms, -2 ms]) and Hilbert transformed to obtain the instantaneous pre-TMS phase of each epoch. In parallel, a second broader window of interest ([-400 ms, -3 ms]) was identified to calculate the power spectra with complex Morlet wavelets. Only individual trials exceeding 50^th^ percentile (>50%max) of power were identified and considered for further analysis. We then classified trials according to their pre-TMS instantaneous phase ([-5 ms, -2 ms]) to retain those at the *peak* (330°-30°) and at the *trough* (150°-210°) of the ongoing oscillation, computed the ERSP and ITPC, and finally Z-transformed the signals ([-0.5 s to 0.6 s]).

The statistical analyses of ERSP and ITPC involved nonparametric (two-tailed dependent samples t-test) cluster-based permutation statistics (5000 iterations controlling for false discovery rate ^68^), comparing each ACTIVE TMS condition to its associated SHAM (e.g., ACTIVE Hi-TMS *vs.* SHAM Hi-TMS / ACTIVE Li-TMS *vs.* SHAM Li-TMS). The statistical analyses of phase-dependent ERSP and ITPC involved nonparametric (two-tailed dependent samples t-test) cluster-based permutation statistics (5000 iterations), comparing each TMS phase-dependent condition (SHAM subtracted) to its associated ACTIVE antiphase (e.g., Hi-TMS Peak (SHAM subtracted) *vs.* Hi-TMS Trough (SHAM subtracted) / Li-TMS Peak (SHAM subtracted) vs. Li-TMS Trough (SHAM subtracted)). When statistical significance criterion was met, post-hoc tests comparing ACTIVE (Peak or Trough) with their respective SHAM conditions (e.g., ACTIVE Hi-TMS SP Peak vs. SHAM Hi-TMS SP Peak / ACTIVE Hi-TMS Rhythmic Trough vs. SHAM Hi-TMS Rhythmic Trough; etc.) were conducted. Monte Carlo p-values were calculated via a two-tailed test (alpha cluster: p<0.05, alpha threshold: p<0.05). The time domain for statistical testing was set from the TMS onset until the epoch end ([0 ms, 600 ms]), and frequency domain centered around the TMS chosen frequency ([8-35 Hz]). Finally, effect sizes (Cohen’s *d*) were computed on the z-score data for any of the identified significant time-frequency clusters.

## Data availability statement

The datasets generated and/or analyzed during the current study are not publicly available due to institutional and ethical restrictions concerning the protection of participant confidentiality and privacy. The data may be made available only for research purposes upon reasonable request to the corresponding author. Requests should include a brief description of the proposed use of the data and will be considered in accordance with the relevant institutional and ethical requirements. Access will be granted where the proposed use is consistent with the participants’ consent and applicable institutional regulations.

## Code availability statement

Codes for data analysis are accessible at the permanent link: https://doi.org/10.5281/zenodo.21894220.

## Authorship contributions

XC-T and AV-C conceived and designed the study with conceptual input by MB, SC, SM, RS, AL and MTC. XC-T and MB conducted the experiments and collected the data. XC-T assisted by MB and AV-C, analyzed the data. XC-T, MB and AV-C prepared the first versions of the manuscript. All authors contributed to critically discuss and read prior versions of this project and approved the final version of the manuscript.

## Acknowledgements

We thank Dr. Cécile Galléa and Dr. Tuomas P. Mutanen for their conceptual contributions and support with data analyses, as well as Ms. Vridhi Rohira and Mr. Carlo Leto for their assistance with data collection.

## Supplementary information

Supplementary information, with 2 information materials sets, 11 figures and 1 table accompanies the main manuscript.

## Supplementary Note 1

For high-intensity blocks, TMS intensity was set at a fixed level of 60% of the maximum stimulator output (MSO). Importantly, we verified that this level of intensity warranted the full recharge of the TMS stimulation boosters and an accurate and consistent delivery of TMS 4 pulse bursts avoiding intensity inaccuracy or missing pulses during rhythmic TMS patterns. Prior evidence has demonstrated that an intensity of ∼80% of the resting motor threshold (rMT) is sufficient to induce measurable scalp EEG responses in absence of peripheral MEPs, hence minimizing the interference of muscle artifacts[1,2] or proprioceptive afferent feedback. Importantly, our post-hoc analyses (see Supplementary Table 1) confirmed that the fixed intensity of 60% MSO used in our study corresponded to sub-motor threshold stimulation (mean relative intensity respect motor threshold = 78,6%rMT), thus minimizing scalp EEG muscular contamination and sensory reafferent feedback after hand muscular evoked contraction.

For low-intensity TMS blocks, we aimed to induce on the M1 surface voltage density levels similar to those applied in rat studies published with the participation of our lab and accounting for currents of ∼0.05-0.1V/m. Given obvious differences between rodent and human scalp thickness, we carried out series of MRI-based tetrahedral Finite Element Model (FEM) simulations of TMS M1 current density distribution on a standard average 152 MNI brain at different TMS intensities with this same equipment (Magstim, Rapid2, 70 mm double coil). On such basis, our quest predicted that the intensity matching most closely the one delivered in prior studies with a custom-made micro-TMS stimulator device (∼0.05-0.1V/m) would need to be of ∼1% MSO, 1.14 dI/dtmax [A/μs]). An in-house induction coil probe attached to a digital oscilloscope (Tektronic 6485, USA), served to estimate the lowest magnetic field output (1-100% MSO) that could be consistently delivered with our human use certified rTMS equipment (Magstim Rapid^2^) and coil (70 mm alpha coil) by measuring on its center the strength of the evoked magnetic field peak voltage at increasing TMS intensities (from 1-10%MSO). We determined that the minimal intensity offering consistent regular pulse output was 3% MSO (3.44 dI/dtmax [A/μs]). Accordingly, 3% MSO (predicted to achieve on M1 a current density of ∼5-12V/m depending on interindividual head volume differences across subjects) was employed in all our participants as the closest proxy of very weak intensity stimulation (compared to the one tested in rodents).

## Supplementary Note 2

To improve the signal-to-noise ratio of our dataset, we implemented a spatial filter that isolated activity emerging from the primary targeted region (the left M1 hotspot for the FDI muscle) and minimized the contribution of multisensory responses (particularly likely evoked during the high-intensity TMS condition at 60%MSO). Since the TMS coil position on the precentral gyrus M1 region differed slightly across participants, this procedure allowed for individualized EEG signal reconstruction. To this end, we first modeled the electrical field (E-field) induced by TMS on each participant’s head and brain volumes. Tetrahedral Finite Element head Models (FEM) were generated from each individual structural T1-weighted MRI scans using the ‘charm’ routine of SimNIBS4.0[3,4]. Electrical fields were then estimated assuming a quasi-static regime[5-7]. TMS coil spatial coordinates [X, Y, Z] for coil simulation were individually extracted from real-world coil position documented during the experimental session with MRI-based neuronavigation software. Then E-field distribution models were computed with a Magstim figure-of-eight 70 mm alpha coil (D70) powered by a MagstimRapid2 (114.7 dI/dtmax [A/μs] = 100% MSO). This included models for each participant for the 3%MSO and 60%MSO TMS intensity conditions employed during our experiments.

The meshes contained 10 different volumes (0.6mm isotropic resolution) segmented from the following tissues: white matter (WM), grey matter (GM), cerebrospinal fluid (CSF), bone, scalp, eyes, compact bone (CB), spongy bone (SB), blood vessels and muscles. The following isotropic conductivity values were assigned: 0.126 S/m (WM), 0.275 S/m (GM), 1.654 S/m (CSF), 0.01 S/m bone, 0.16 S/m scalp, 0.5 S/m eyes, 0.008 S/m CB, 0.025 S/m SB, 0.6 S/m blood vessels, and 0.46 S/m muscles. Tissue segmentation was performed automatically relying on CAT12 and SPM12 and manually corrected when required after careful visual inspection. Electrical fields were estimated assuming a quasi-static regime, according to

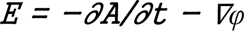

where E is the electric field vector and φ the electric potential. Models were computed with a Magstim figure-of-eight 70 mm coil (D70) powered by a MagstimRapid^2^ (114.7 dI/dtmax [A/μs] = 100%MSO). Since the relationship between A/μs and %MSO is linear, TMS coil rate of change (dI/dt) was set to 3.4A/μs for the low intensity TMS simulations (Li-TMS, 3% MSO) and to 68.8A/μs for high-intensity models (Hi-TMS, 60% MSO). TMS coils were spatially placed according to the neuronavigation actual positions (Brainsight, Rogue Solutions) used during the experimental session and manually corrected when necessary, maintaining a 7mm coil-to-scalp distance to simulate the presence of active EEG electrodes. Biophysical simulations of current distribution in our study served two purposes. Firstly, they allowed us to estimate for each individual participant the peak current generated by Li-TMS (3%MSO) and Hi-TMS (60%MSO) intensities employed on the left M1 cortical target. Secondly, they served to construct individual spatial filters to process EEG data at the source level.

Based on the modeled current field distribution, we built an individual spatial filter with special weighting constraints fulfilled by E-field outcomes. The procedure was based on a previously developed framework for the design of crosstalk functions for spatial filtering applied to EEG/MEG data (DeFleCT)[8]. The implemented procedure employed the tools SimNIBS 4.0, ISO2MESH[9], the Helsinki BEM framework[10], EEGLAB, TESA toolbox, DeFleCT framework and operated as follows: From the E-field simulations we first outlined the primary impacted cortical structure (>80% of the maximal TMS-induced E-field strength). In parallel, we computed the leadfield matrix from the derived forward models. Finally, the cross-talk leakage of our sources of interest (sources within the magnetic field boundaries (>80%)) was estimated and used as a source-based spatial filter (aka cross-talk function) employing a minimum norm like estimator for every participant and applied to the EEG data delimiting pass-band sources and minimizing cross-talk leakage from other non-highlighted sources anywhere in the brain. The procedure source-informed, reconstructed and separated our EEG signals of interest (i.e., from the primarily targeted cortical source) while minimizing crosstalk from other locations anywhere else in the brain. Details of E-field simulations, ROIs and crosstalk leakage of the employed spatial filters can be found in Supplementary Figures.

## Supplementary References

**Supplementary Figure 1.**
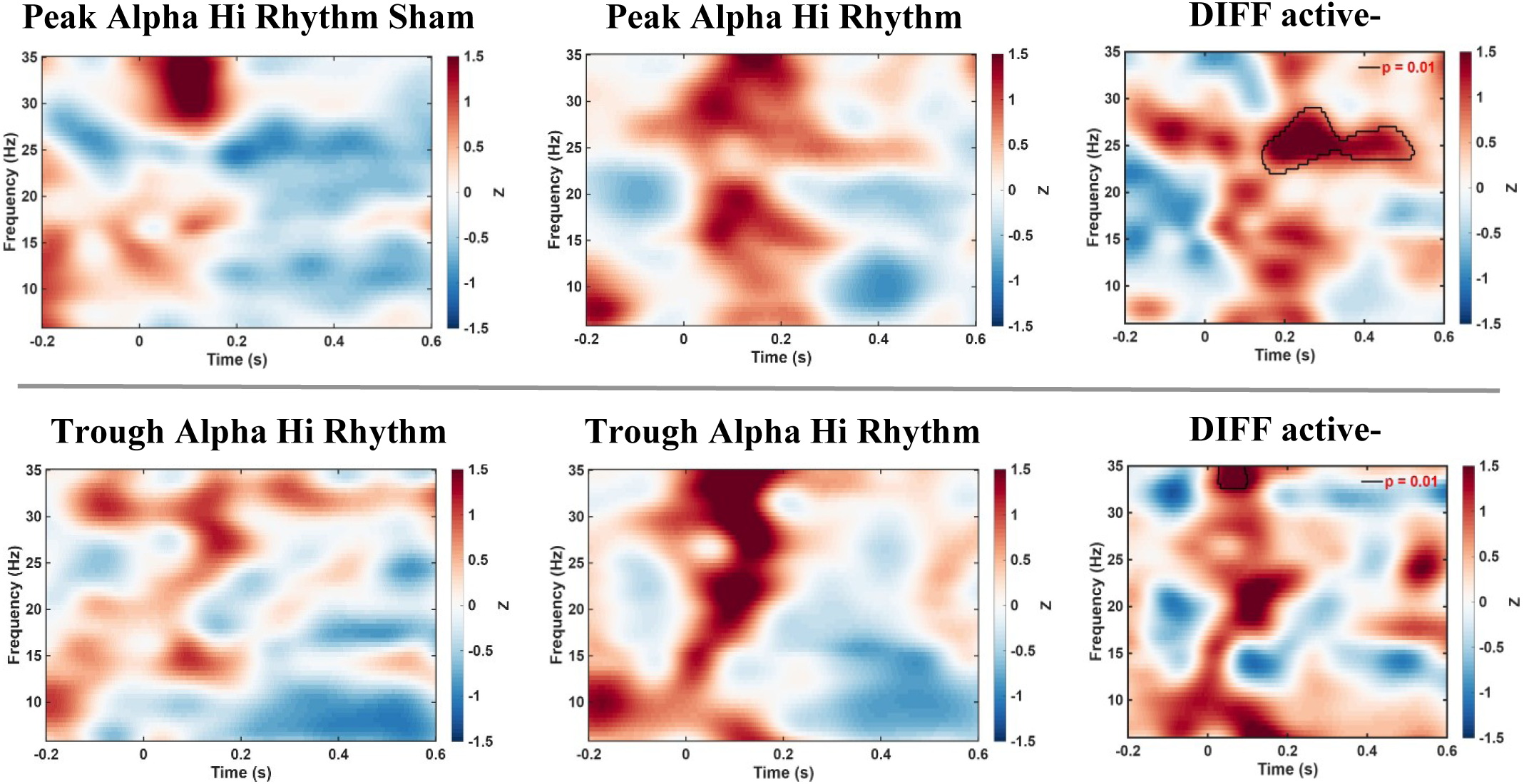
“Post Hoc” active *vs.* sham comparisons for Hi-Rhythmic Alpha pre-TMS phase dependent ERSP. **Effects of high-intensity rhythmic TMS on event-related spectral perturbation (ERSP).** The upper panels shows the ERSP for trials in which TMS was delivered at the peak of the ongoing alpha rhythm, whereas the lower panel shows the ERSP for trials in which stimulation was delivered at the trough of the ongoing alpha rhythm. For each panel, sham (left), active (middle), and the statistical comparison between active and sham conditions (right) are displayed. The ERSP maps represent *z*-transformed grand-average data across participants for each TMS condition. Black contours indicate statistically significant clusters in the time–frequency domain (*p* < 0.05, two-sided).

**Supplementary Figure 2.**
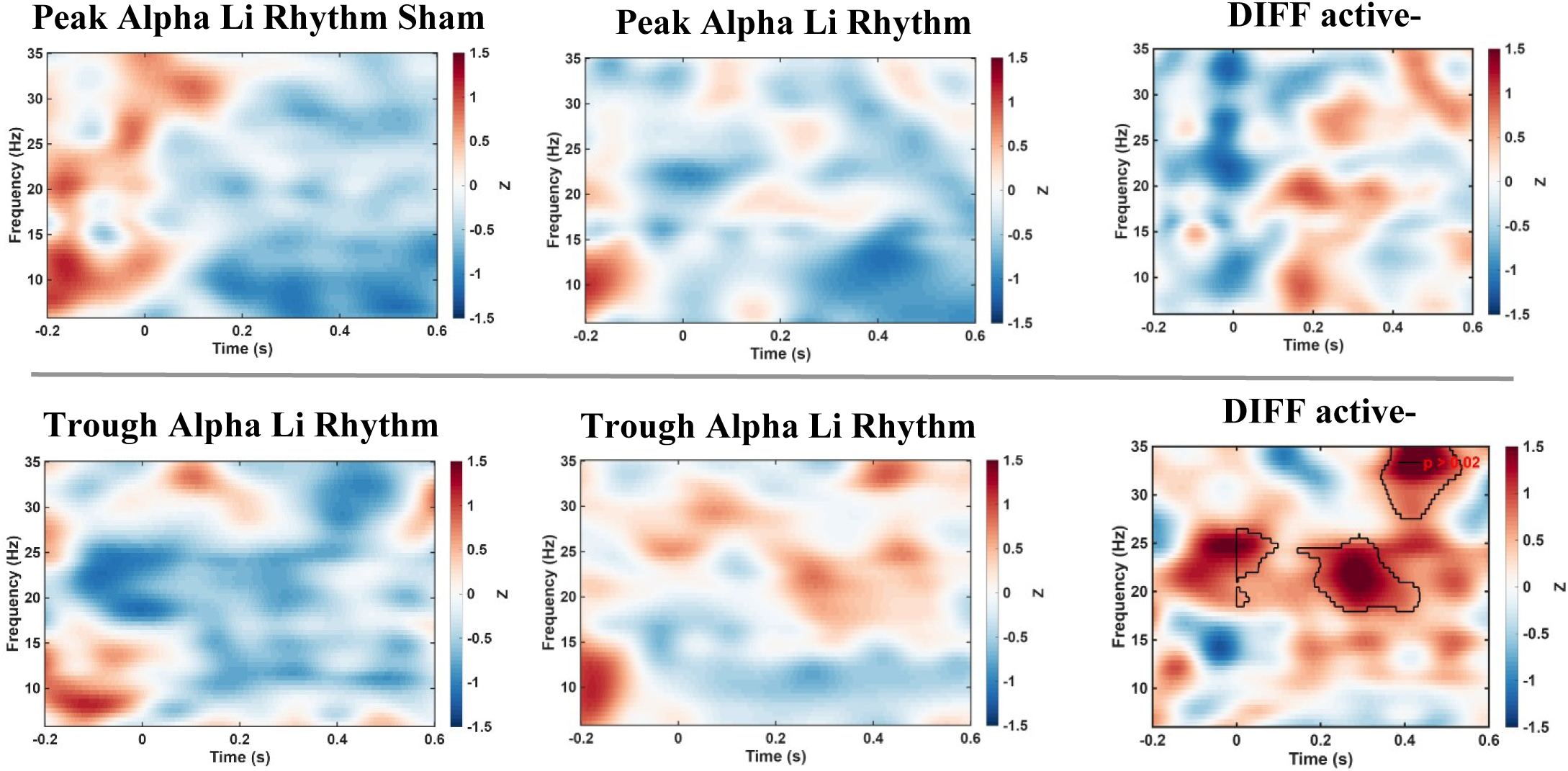
“Post Hoc” active *vs.* sham comparisons for Li-Rhythmic Alpha pre-TMS phase dependent ERSP. **Effects of low-intensity rhythmic TMS on event-related spectral perturbation (ERSP)**. The upper panels shows the ERSP for trials in which TMS was delivered at the peak of the ongoing alpha rhythm, whereas the lower panel shows the ERSP for trials in which stimulation was delivered at the trough of the ongoing alpha rhythm. For each panel, sham (left), active (middle), and the statistical comparison between active and sham conditions (right) are displayed. The ERSP maps represent *z*-transformed grand-average data across participants for each TMS condition. Black contours indicate statistically significant clusters in the time–frequency domain (*p* < 0.05, two-sided).

**Supplementary Figure 3.**
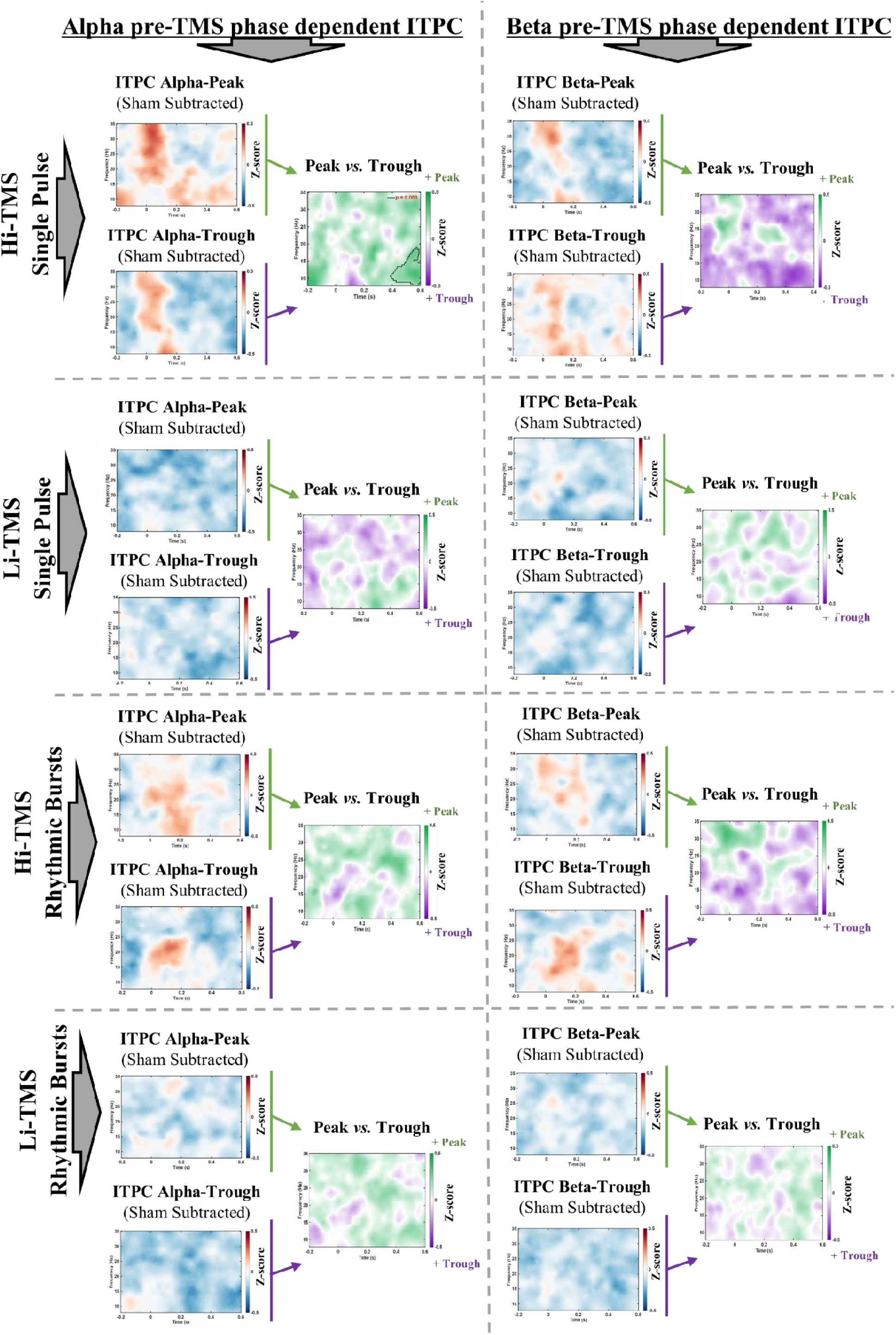
Phase dependent ITPC. **Alpha and Beta phase-dependent TMS effects on Inter-trial Phase Coherence (ITPC) estimates.** Group-level (n=18) ITPC measurements sorted according to the instantaneous pre-TMS onset phase (-5ms to -2ms) of the mu-alpha rhythm (8-13Hz, left column) and beta rhythm (14-30Hz, right column). ACTIVE TMS trials (exceeding +50% of total ERSP oscillatory power (-400ms to -3ms) across all ACTIVE trials, SHAM subtracted) at the *peak* or the *trough* of the alpha and beta rhythm at the time of 1st TMS pulse onset were identified and selected for final analysis. Per every tested TMS condition, the phase-dependent ITPC time-frequency Z-score for ACTIVE (SHAM subtracted) is independently displayed, where ***Red areas*** indicate ITPC increase, and ***Blue areas*** indicate ITPC decrease. Moreover, the comparisons between TMS conditions contrasting *peak* vs *trough* sorted trials are displayed in the figure. ***Green clusters*** of EEG plots indicate ITPC increase when TMS pulses were delivered around the *peak* of the ongoing oscillations. ***Purple clusters*** of EEG plots indicate ITPC increase when TMS pulses were delivered around the *trough* of the ongoing oscillations. Black outlines indicate statistically significant clusters between conditions in the time and frequency domains (p<0.05, two-sided).

**Supplementary Figure 4.**
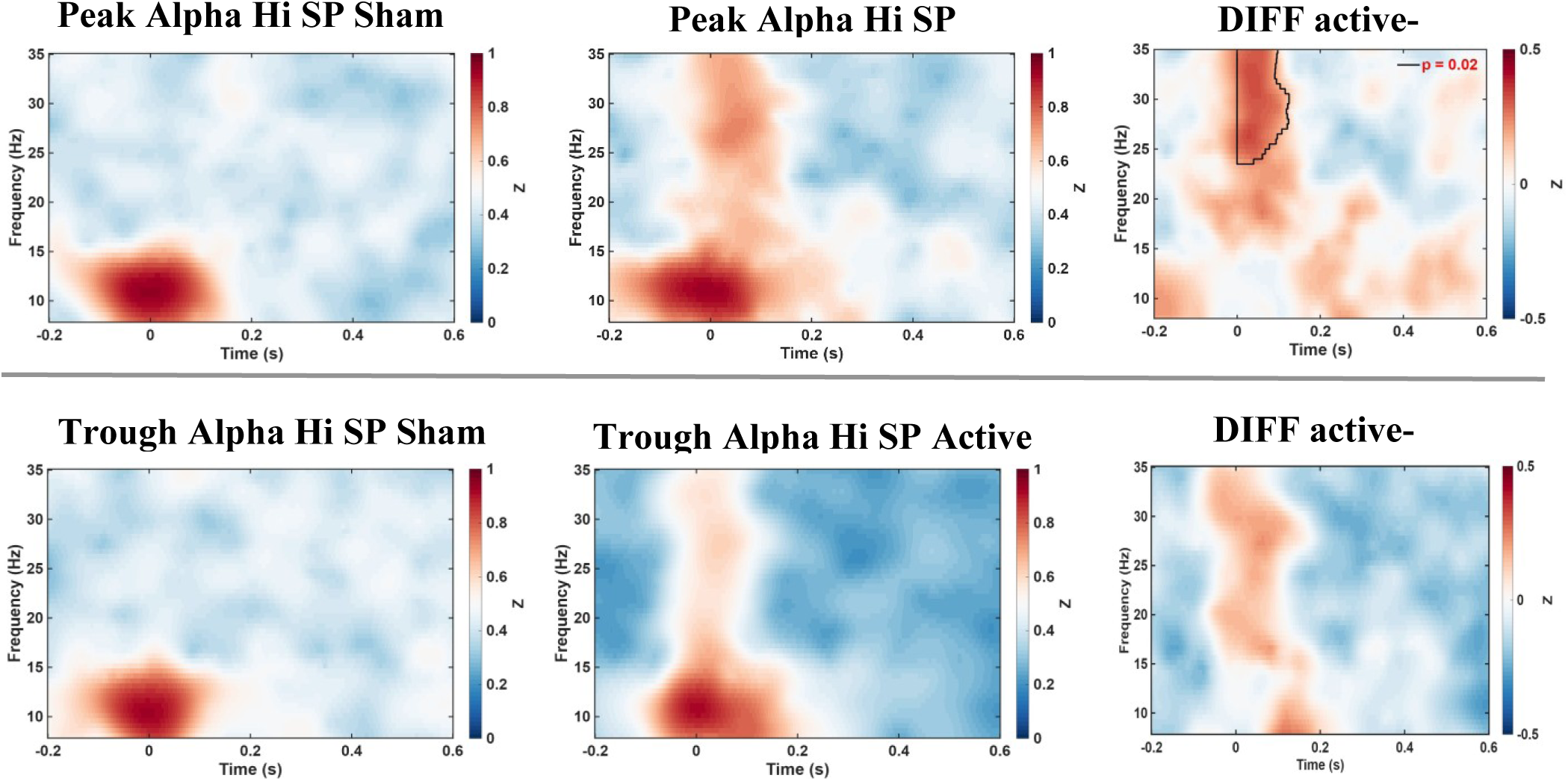
“Post Hoc” active *vs.* sham comparisons for Hi-Single Pulse Alpha pre-TMS phase dependent ITPC. **Effects of high-intensity single pulse TMS on inter trial phase coherence (ITPC).** The upper panels shows the ITPC for trials in which TMS was delivered at the peak of the ongoing alpha rhythm, whereas the lower panel shows the ITPC for trials in which stimulation was delivered at the trough of the ongoing alpha rhythm. For each panel, sham (left), active (middle), and the statistical comparison between active and sham conditions (right) are displayed. The ITPC maps represent *z*-transformed grand-average data across participants for each TMS condition. Black contours indicate statistically significant clusters in the time–frequency domain (*p* < 0.05, two-sided).

**Supplementary Figure 5.**
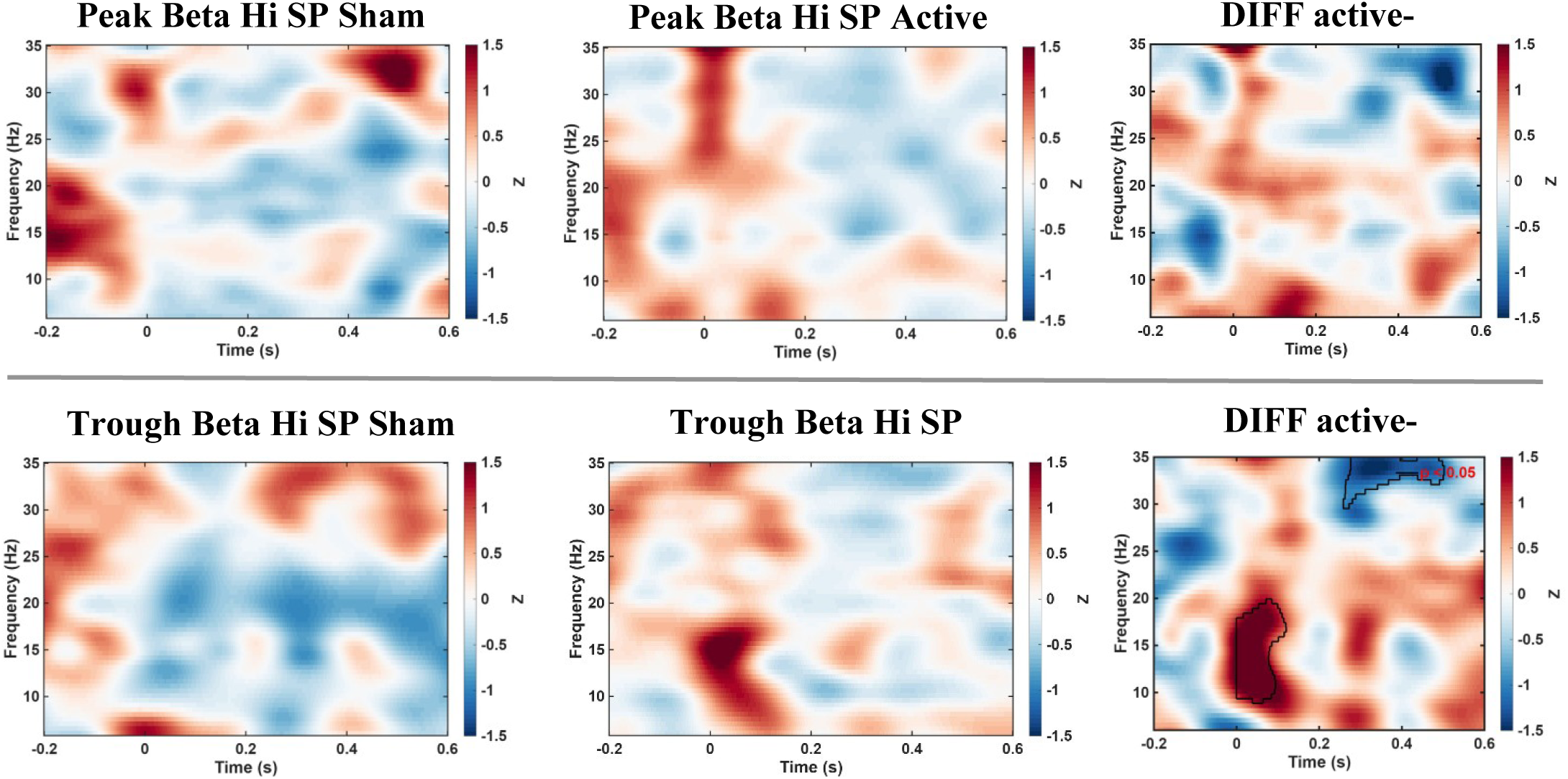
“Post Hoc” active *vs.* sham comparisons for Hi-Single Pulse Beta pre-TMS phase dependent ERSP. **Effects of high-intensity single pulse TMS on event-related spectral perturbation (ERSP).** The upper panels shows the ERSP for trials in which TMS was delivered at the peak of the ongoing beta rhythm, whereas the lower panel shows the ERSP for trials in which stimulation was delivered at the trough of the ongoing beta rhythm. For each panel, sham (left), active (middle), and the statistical comparison between active and sham conditions (right) are displayed. The ERSP maps represent *z*-transformed grand-average data across participants for each TMS condition. Black contours indicate statistically significant clusters in the time–frequency domain (*p* < 0.05, two-sided).

**Supplementary Figure 6.**
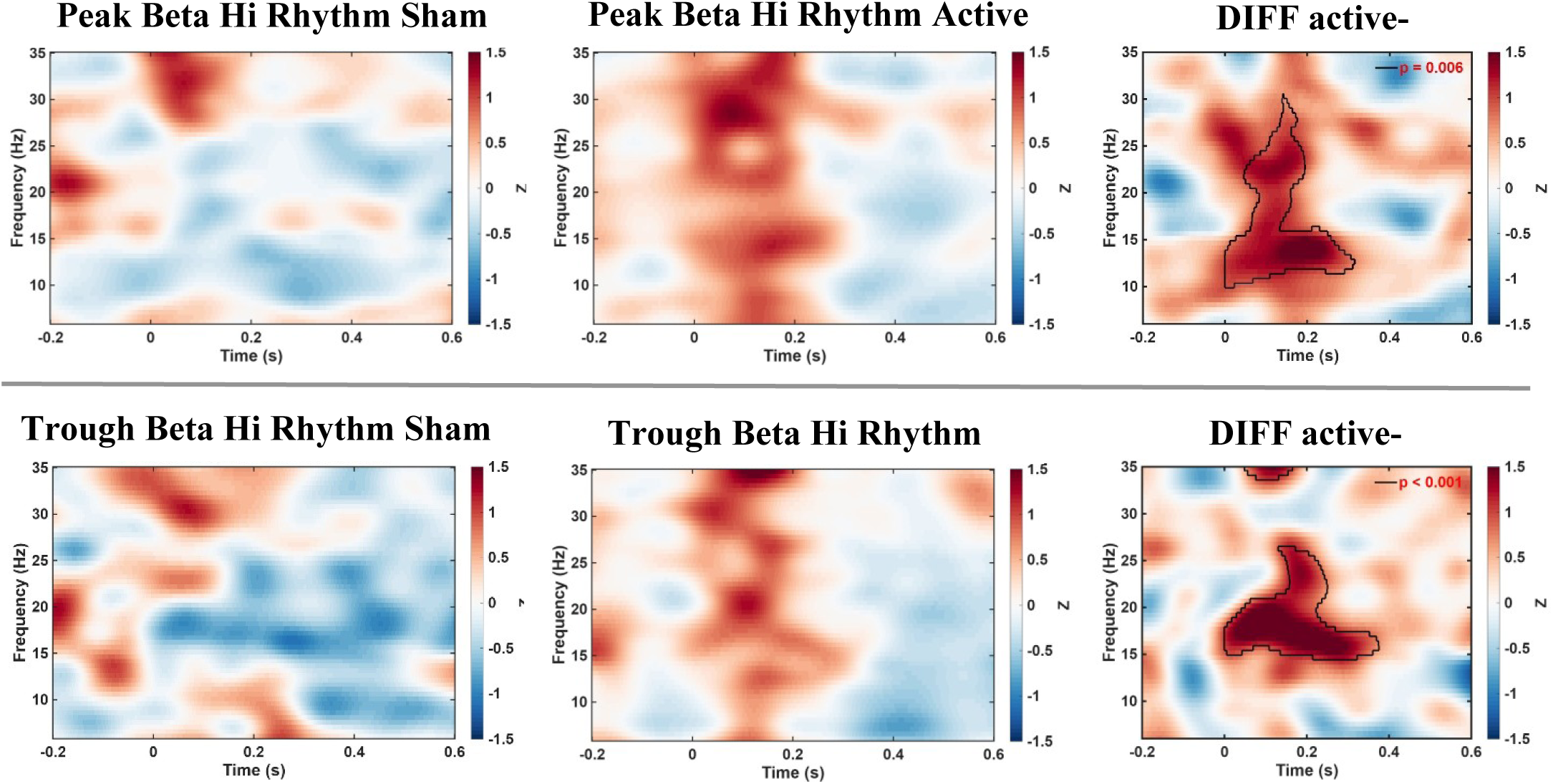
“Post Hoc” active *vs.* sham comparisons for Hi-Rhythmic Beta pre-TMS phase dependent ERSP. **Effects of high-intensity rhythmic TMS on event-related spectral perturbation (ERSP).** The upper panels shows the ERSP for trials in which TMS was delivered at the peak of the ongoing beta rhythm, whereas the lower panel shows the ERSP for trials in which stimulation was delivered at the trough of the ongoing beta rhythm. For each panel, sham (left), active (middle), and the statistical comparison between active and sham conditions (right) are displayed. The ERSP maps represent *z*-transformed grand-average data across participants for each TMS condition. Black contours indicate statistically significant clusters in the time–frequency domain (*p* < 0.05, two-sided).

**Supplementary Figure 7.**
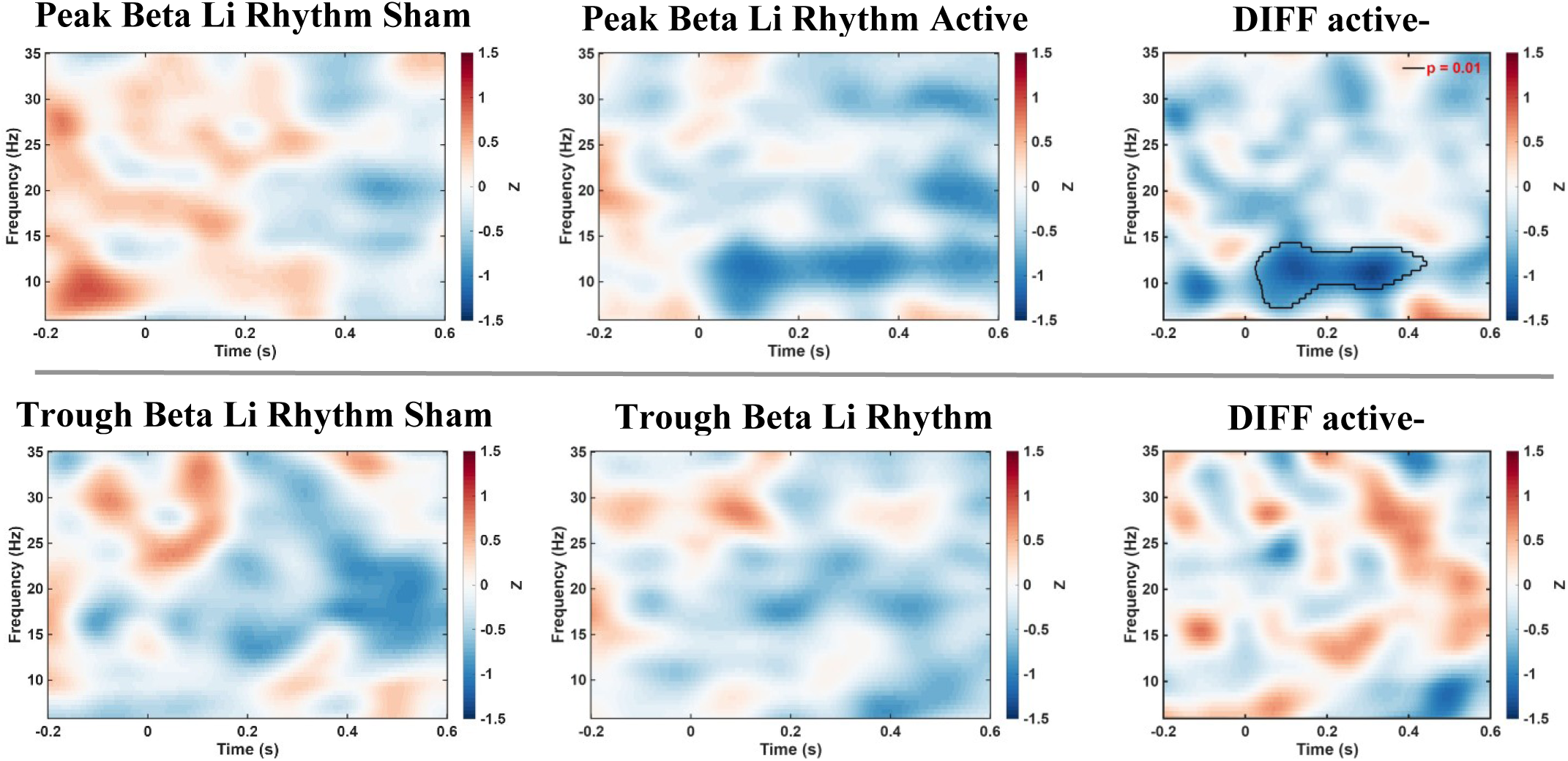
“Post Hoc” ERSP active *vs.* sham comparisons for Li-Rhythmic Beta pre-TMS phase dependent. **ERSP Effects of low-intensity rhythmic TMS on event-related spectral perturbation (ERSP).** The upper panels shows the ERSP for trials in which TMS was delivered at the peak of the ongoing beta rhythm, whereas the lower panel shows the ERSP for trials in which stimulation was delivered at the trough of the ongoing beta rhythm. For each panel, sham (left), active (middle), and the statistical comparison between active and sham conditions (right) are displayed. The ERSP maps represent *z*-transformed grand-average data across participants for each TMS condition. Black contours indicate statistically significant clusters in the time–frequency domain (*p* < 0.05, two-sided).

**Supplementary Figure 8.**
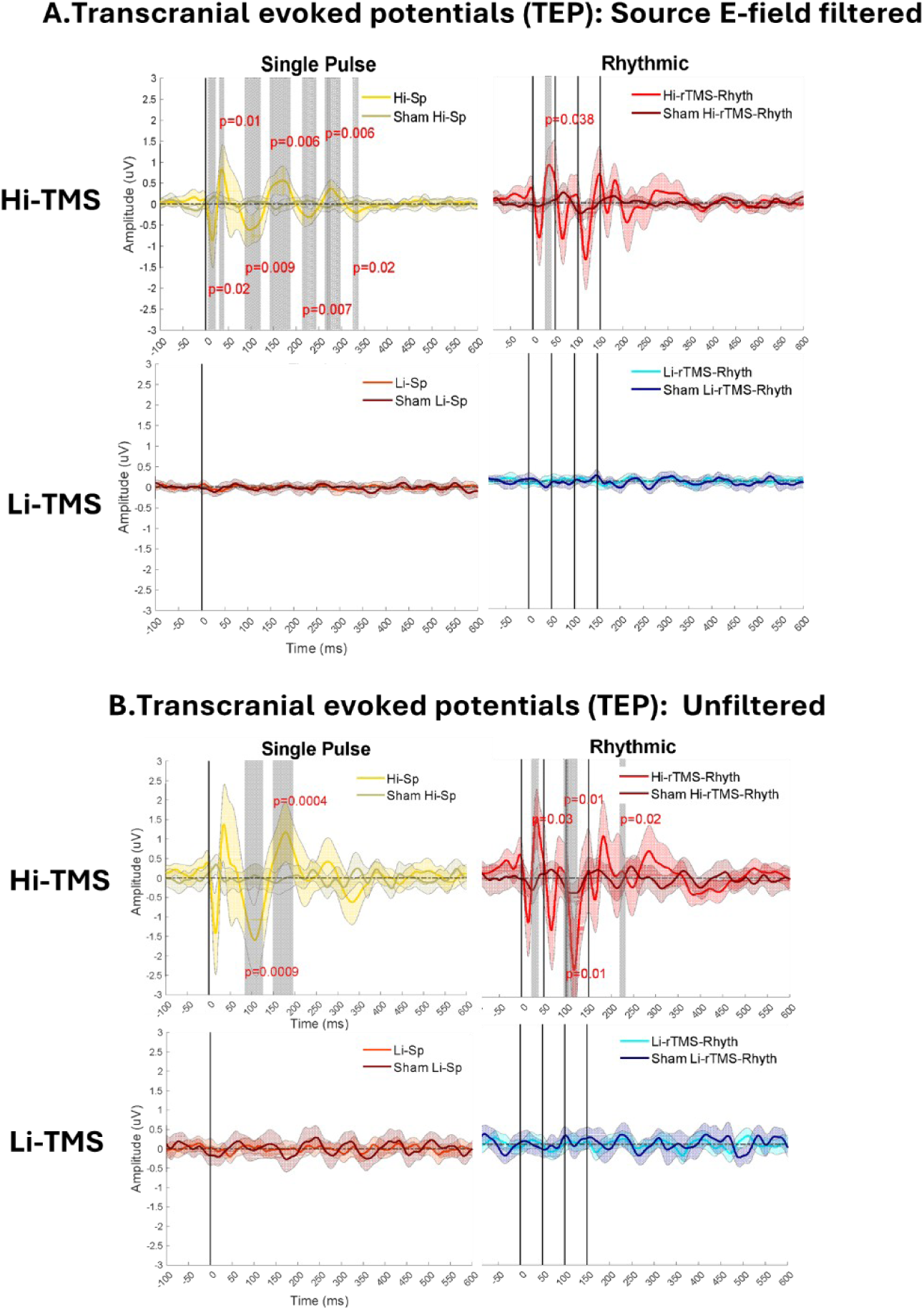
Transcranial Evoked Potentials (TEPs) **Time-course of Transcranial Evoked potentials (TEPs)**. All trials have been averaged across participants and grouped by specific TMS conditions at high (Hi-TMS), and at very weak-intensity (Li-TMS) stimulation. The group mean and standard deviation (±SD) values for each condition are indicated with thick and a thin line profiles, respectively. Vertical straight solid lines signal the onset time of each individual TMS pulse. Vertical grey-shaded areas indicate statistically significant EEG clusters in the time and frequency domains (Montecarlo cluster based permutation statistics, cluster alpha p<0.05, Montecarlo alpha p<0.05, two-sided) across conditions. Hi-TMS: high-intensity TMS (60%MSO, ∼113V/m), Li-TMS: low-intensity (3%MSO, ∼6V/m). **A.** Spatially filtered TEP for all conditions. **B.** TEP responses of non-spatially filtered signals (original EEG signals excluding the application of E-field-informed spatial filtering procedures implemented in our final analyses) at the group level from a cluster of electrodes overlying the stimulated left primary motor (M1). The EEG electrodes selected for the analyses were: C3, C5, CP1, C1 and CP3. For this figure, TEP analyses and statistical comparisons were applied as described in the main manuscript but excluding spatial filtering preprocessing step in sensor level TEP time courses.

**Supplementary Figure 9.**
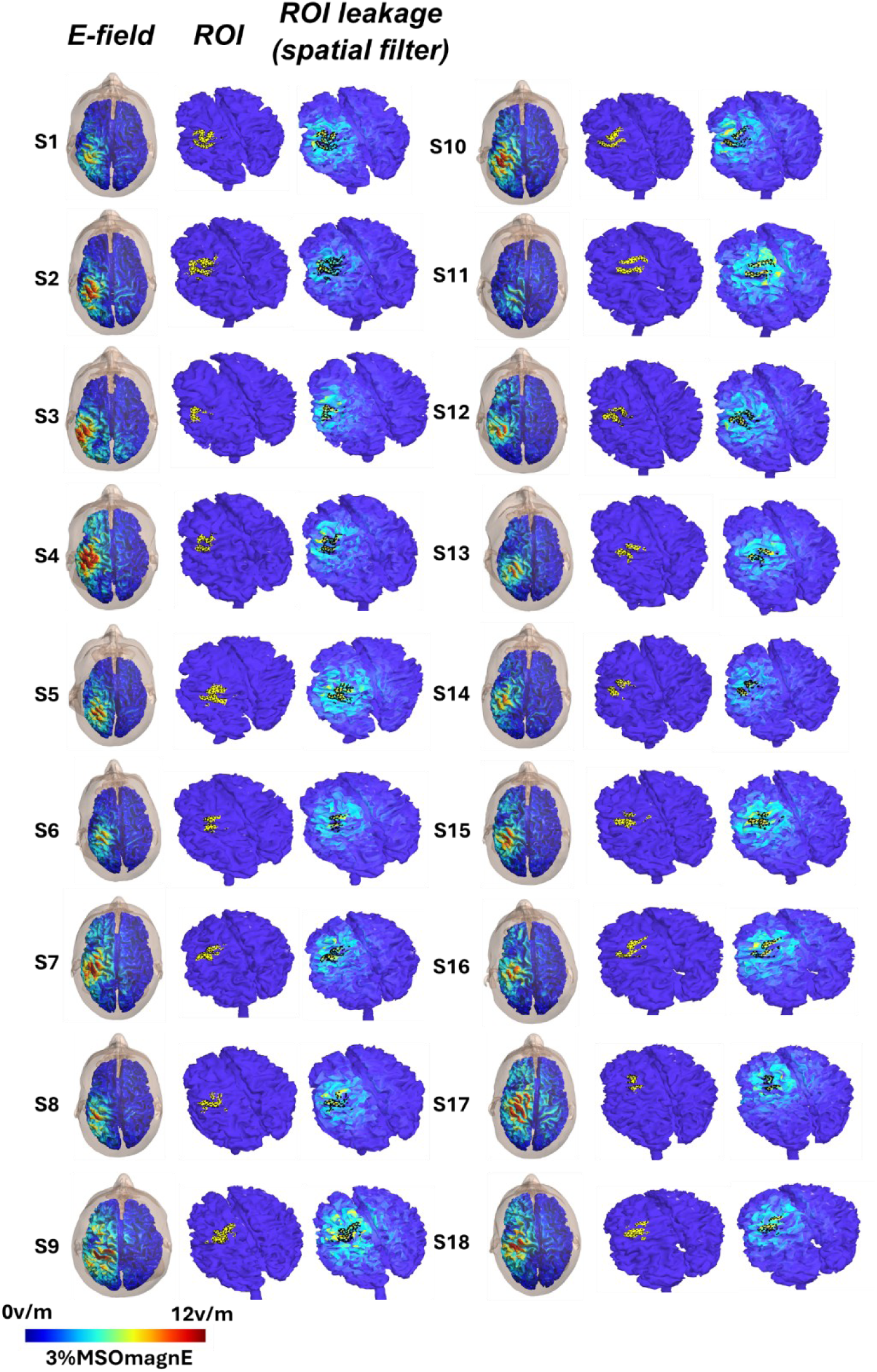
E-field models and spatial filter. **Electric field modeling and spatial filters.** MRI-based E-field spread simulations were used to define the “boundaries” of the estimated region of interest (ROI) primarily affected by the TMS pulses or bursts. For each participant (S1–S18), the ROI was determined using the 80th percentile of the maximum peak field strength and the spatial filter’s crosstalk leakage sensitivity. To aid visualization, all individual E-field values were normalized on a scale from 0 V/m (minimum) to 12 V/m (maximum). The simulations were generated by reconstructing each participant’s structural head model using SIMNIBS 4.0. In the ROI leakage figures, any sources outside the highlighted ROI were minimized through the spatial filtering procedure. As a result, the crosstalk leakage maps can also be interpreted as participant-specific reconstructions of the filtered signal source location.

**Supplementary Figure 10.**
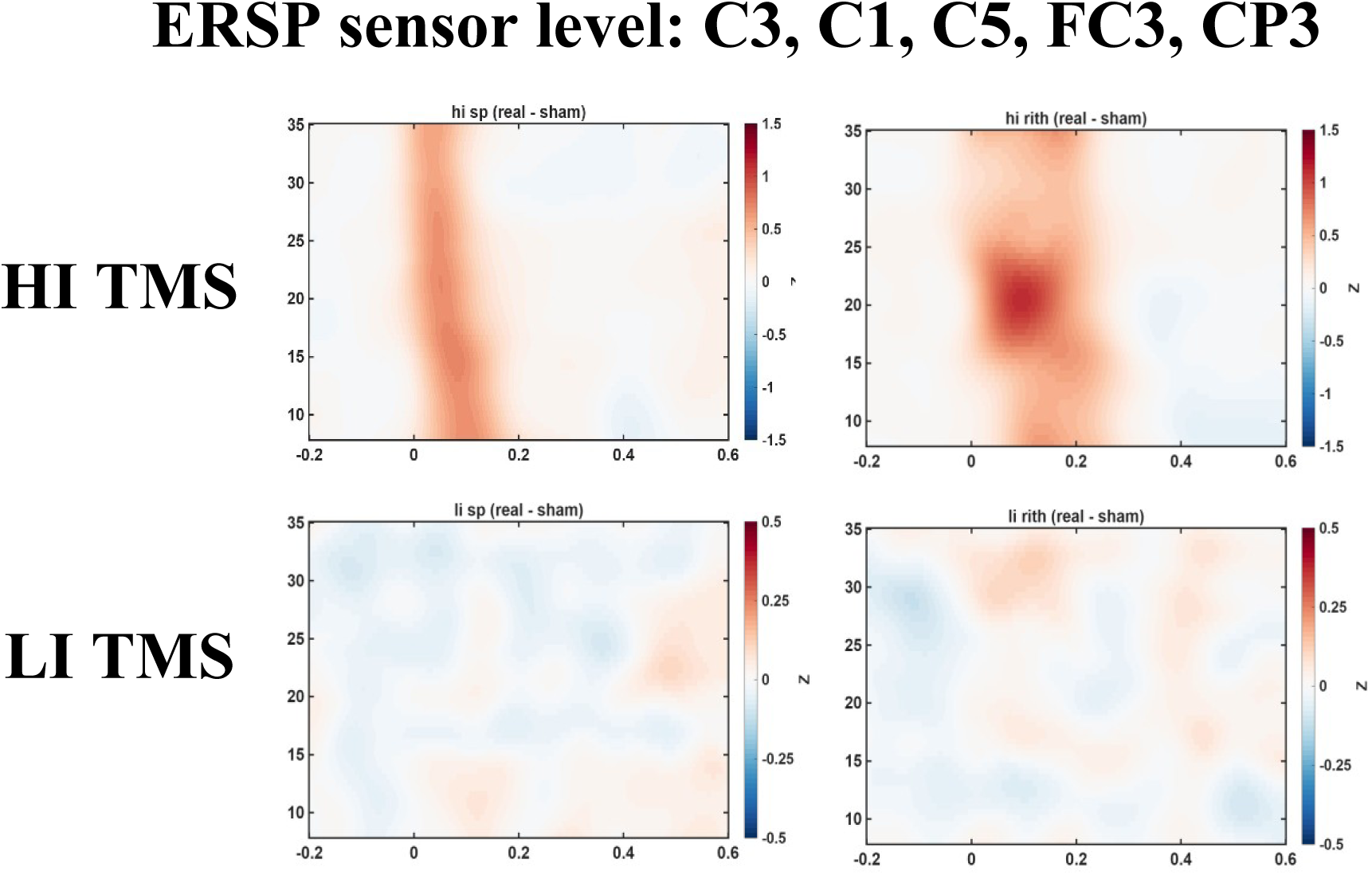
ERSP obtained from original sensor level data (non-spatially filtered) **ERSP time-frequency maps of non-spatially filtered EEG signals** (i.e., original EEG signals analyzed without applying the E-field-informed spatial filtering procedure implemented in the primary analyses). Group-level ERSP were calculated from a cluster of electrodes overlying the stimulated left primary motor cortex (M1), comprising electrodes C3, C5, CP1, C1, and CP3.

**Supplementary Figure 11.**
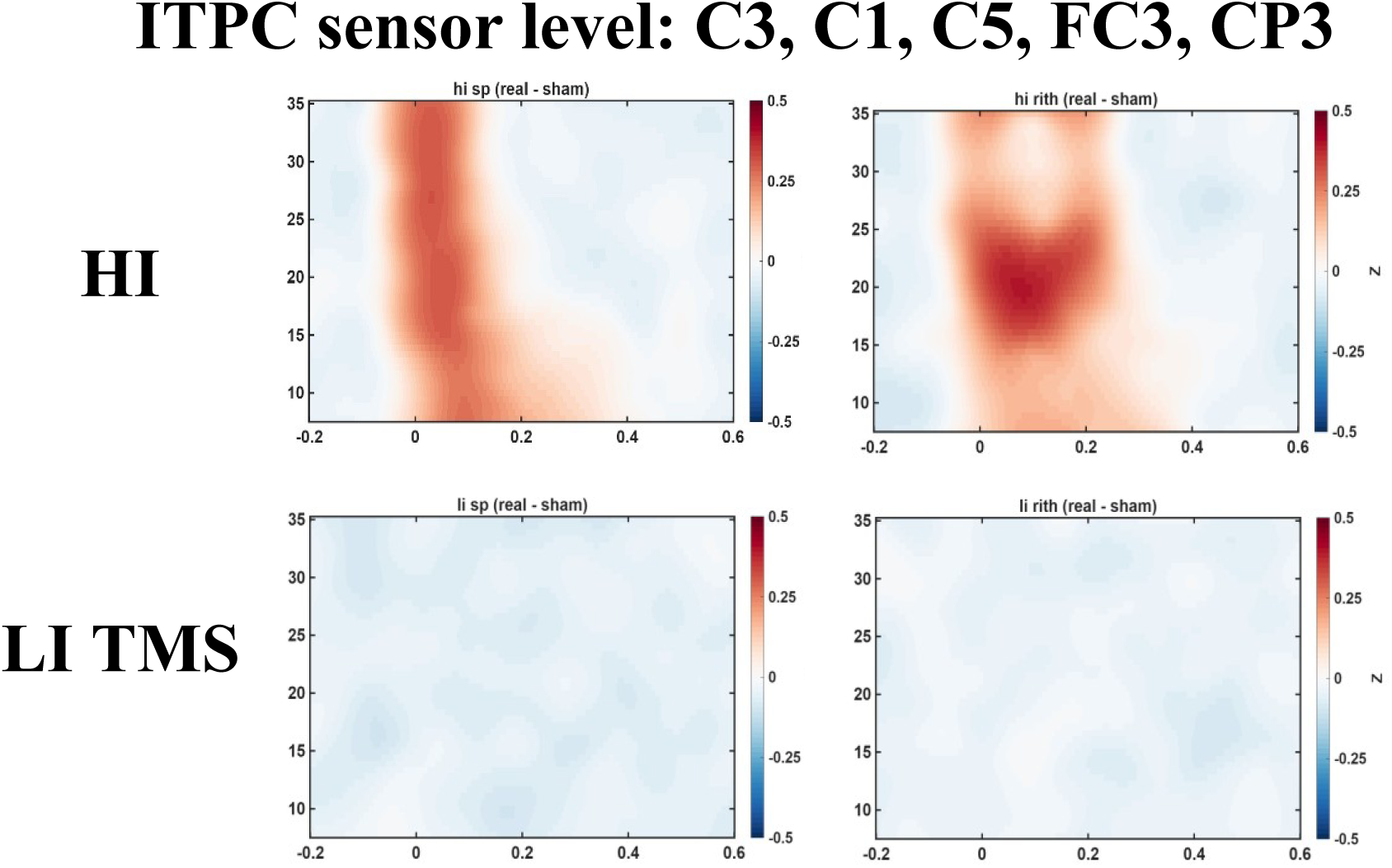
ITPC obtained from original sensor level data (non-spatially filtered) **ITPC time-frequency maps of non-spatially filtered EEG signals** (i.e., original EEG signals analyzed without applying the E-field-informed spatial filtering procedure implemented in the primary analyses). Group-level ITPC were calculated from a cluster of electrodes overlying the stimulated left primary motor cortex (M1), comprising electrodes C3, C5, CP1, C1, and CP3.

**Supplementary Table 1.**
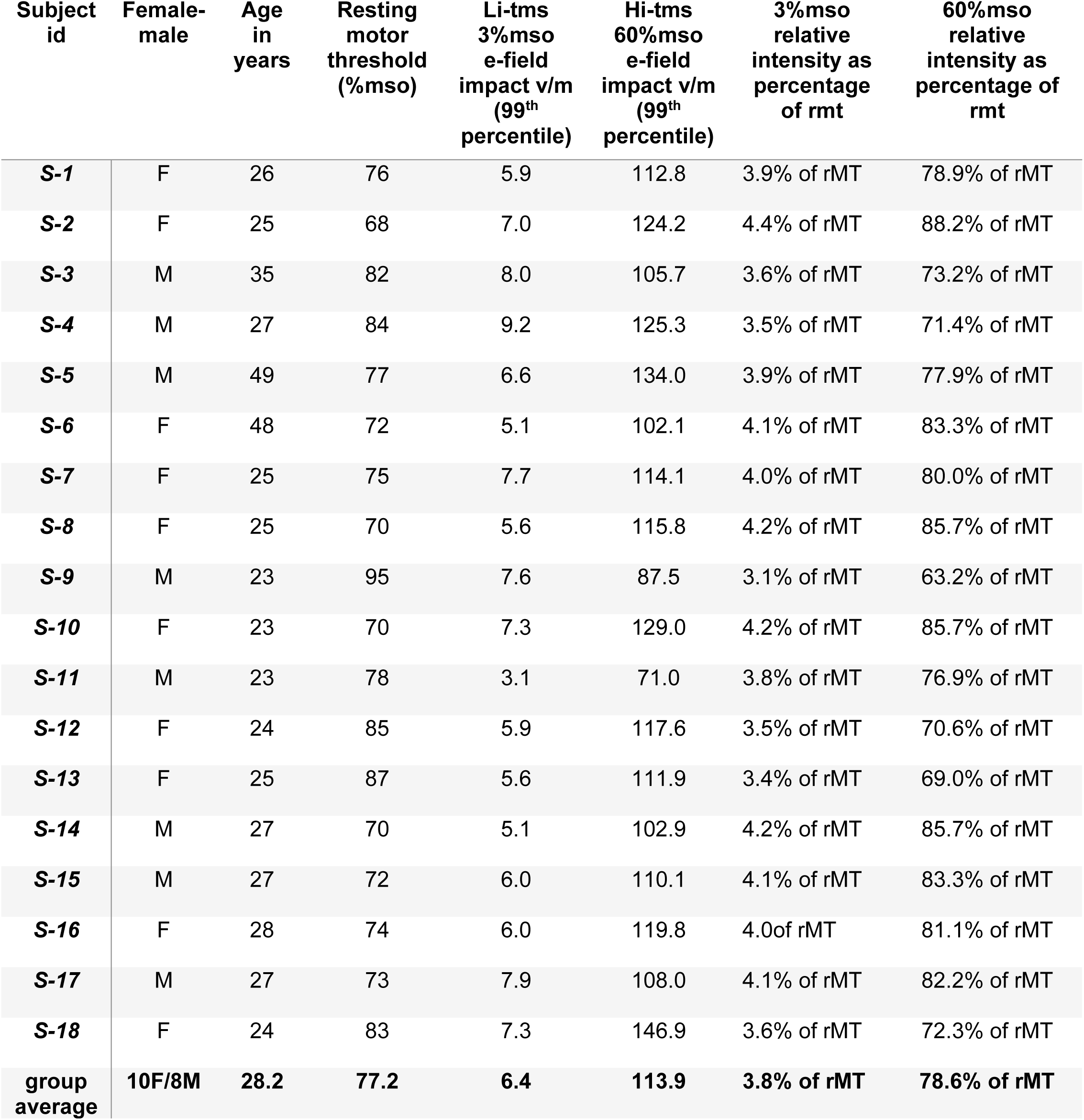
Demographic information. **Demographic characterization of the study cohort.** Sex (F: Female, M: Male), Age (in years), resting motor threshold (% Maximal Stimulator’s output, MSO), individual values of the 99^th^ percentile E-field peak value (V/m) extracted from MRI-based current field models and simulations estimated on the grey matter, and relative motor threshold at fixed MSO employed during the experiment. Prior to the start of the session, we measured and documented for each participant their resting Motor Threshold (rMT). To this end, TMS single pulses were delivered over the M1 hotspot corresponding to the right FDI hand muscle while we recorded the amplitude of the MEPs (Signal 5.0, Cambridge Electronic Design Limited, Cambridge, UK). Starting at 65%MSO, TMS was delivered at varying intervals between ∼6-8 secs, with intensity increases/decreases by steps of 1-5% of the MSO, depending on the consistency and amplitude of evoked MEPs responses. The individual rMT was defined as the lowest TMS intensity able to induce at least 5 out of 10 MEP (50% responses) with a minimal peak-to-peak amplitude of 50μV. Note that resting motor threshold was estimated with EEG cap mounted.

